# 3-Phenylpropionic acid, a microbiota-derived polyphenol metabolite linked to hippuric acid, ameliorates colitis as a colon-enriched PPAR-γ agonist

**DOI:** 10.1101/2025.11.25.690607

**Authors:** Liyuan Xiang, Wanrong Luo, Wenjie Zhang, Wenhua Lai, Xueting Wu, Shuyu Zhuo, Boya Feng, Sifan Chen, Shuguang Fang, Minhu Chen, Peng Bai, Rui Feng, Yijun Zhu

## Abstract

Polyphenols are promising therapeutics for Crohn’s disease (CD), yet their bioavailability is largely dependent on the gut microbiome. The bioactive metabolites, metabolic pathways, and anti-inflammatory mechanisms underlying their effects remain poorly defined. We identified hippuric acid (HIPA) as a biomarker of polyphenol–microbiome interaction. HIPA is a meta-organismal metabolite produced through a cross-organ pathway that involves synthesis from gut microbial polyphenol metabolism *via* 3-phenylpropionic acid (3-PPA), followed by renal clearance. We found that CD is characterised by impaired polyphenol metabolism, resulting in deficient colonic 3-PPA and reduced serum hippuric acid relative to healthy individuals. Furthermore, we demonstrate that 3-PPA is a colon-enriched, mild PPAR-γ agonist that ameliorates colitis in murine model. Systemic bioinformatic profiling of metagenomic data suggested the existence of undiscovered 3-PPA producers. Through targeted microbial incubation with polyphenols, we identified the probiotic *Bifidobacterium breve* and a lab isolate *Escherichia coli* FAH as novel 3-PPA producers. Collectively, our findings establish 3-PPA as a diet-dependent, colon-produced, mild PPAR-γ agonist with the potential to minimise adverse effects associated with traditional PPAR-γ agonists. This work establishes a foundation for developing microbiota-targeted, polyphenol-based therapeutic strategies for CD patients.

---

Crohn’s disease (CD) and ulcerative colitis (UC) are the two primary forms of inflammatory bowel disease (IBD), characterised by symptoms such as diarrhea, rectal bleeding, abdominal pain, fatigue, and weight loss, all of which significantly impair quality of life[1]. The global incidence and prevalence of IBD are rising, particularly in newly industrialised countries, highlighting the urgent need for improved preventive and therapeutic strategies. While the precise etiology of IBD remains unclear, it is widely accepted that disease onset involves complex interactions among environmental factors, host genetics, gut microbiota, and the immune system [1,2]. While dietary intake shapes gut microbial ecology, the metabolic output of this microbiota determines the fate of dietary components and, in turn, influences host immunity, ultimately contributing to intestinal inflammation and the development of CD [3,4].

A plant-based diet, rich in fruits, vegetables, and whole grains, and low in processed meats and refined carbohydrates, has been associated with a reduced risk of CD flare-ups [5,6]. The protective effects of such diets are partially attributed to polyphenols, a diverse class of plant-derived secondary metabolites found in a variety of foods including fruits, vegetables, cereals, olives, legumes, chocolate, tea, coffee, and wine [7]. Polyphenols and their derived phenolic acids hold potential as colon-targeted therapeutic agents. It is estimated that 5–10% of ingested polyphenols are absorbed in the small intestine, while the majority (90–95%) reach the colon, where they can accumulate at millimolar concentrations and are extensively metabolised by the gut microbiota to produce small phenolic acids [8]. Phenolic acids, including 3-phenylpropionic acid (3-PPA, also known as hydrocinnamic acid) and phenylacetic acid, have been detected at submillimolar concentrations in human fecal water, and their levels increase significantly following raspberry supplementation, suggesting that microbial polyphenol metabolism in the colon can generate therapeutically relevant concentrations of bioactive compounds [9,10].

Peroxisome proliferator-activated receptors (PPARs) are nuclear receptor transcription factors activated by endogenous ligands [11]. Among them, PPAR-γ is highly expressed in colonic tissues and have emerged as promising therapeutic targets in IBD due to its roles in regulating inflammation, metabolism, and cellular differentiation [12–14]. The commonly used IBD drug 5-aminosalicylic acid (5-ASA) has been shown to activate PPAR-γ signaling in the colon, contributing to its anti-inflammatory properties [15]. While PPAR-γ agonists such as thiazolidinediones (for example rosiglitazone and pioglitazone) demonstrate efficacy in preclinical models, their clinical utility is limited by serious adverse effects including weight gain, edema, heart failure, and increased fracture risk [16–18], which result from systemic overactivation of PPAR-γ by full agonists [19,20]. These limitations underscore the urgent need for tissue selective partial PPARγ agonists. In this context, microbiota derived colon restricted metabolites highlight a promising therapeutic avenue for harnessing PPAR-γ signaling to attenuate colitis. Indeed, microbiome derived butyrate activates PPAR-γ, enhancing β-oxidation in colonocytes and reducing luminal oxygen availability, although the precise mechanism remains unclear [21].

Significant interindividual variation in gut microbiota composition drives differential metabolism of dietary polyphenols, ultimately contributing to their variable health benefits across populations [22–24]. Our previous study demonstrated that flavonoid degradation, one of the best-characterised polyphenol pathways, has been shown to be diminished in CD patients, as indicated by reduced abundance of flavonoid-degrading genes in their microbiota [25]. To date, only a limited number of microbial polyphenol metabolites and metabolic pathways have been characterised [25,26]. A deeper understanding of polyphenol metabolism fate and mechanism are crucial for illuminating and developing strategies to restore diet-microbiome–host immune homeostasis.

In this study, we aimed to elucidate the metabolic fate of dietary polyphenols by identifying metabolites produced by the gut microbiome and evaluating their potential for personalised diagnostics and precision nutrition in CD. We identified hippuric acid (HIPA) as a biomarker of dietary polyphenols-microbiota interaction. Its microbial precursor, 3-PPA, originates from dietary polyphenols and functions as a mild PPAR-γ agonist that alleviates colitis. Our systematic analysis revealed a broader ecological distribution of microbial 3-PPA biosynthesis. Beyond the established role of *Clostridium sporogenes*, we identified probiotic *Bifidobacterium breve* and a novel isolate, *Escherichia coli* FAH, reduce trans-cinnamic acid to 3-PPA. These findings provide a basis for developing microbiome-targeted precision nutrition utilise dietary polyphenols for the remission management of CD.

## Results

### Elevated HIPA in healthy individuals (HCs) originates primarily from dietary polyphenols

Previous studies have demonstrated considerable interindividual variability in gut microbial polyphenol metabolism, with HCs exhibiting a higher metabolic potential compared to those with CD subjects implied by metagenomics analysis [22,25]. Our primary objective was therefore to identify a serum biomarker that reflects microbial polyphenol metabolic efficiency. We hypothesised that such a biomarker would: 1) contain a benzene ring, 2) be microbiome-derived, 3) be technically convenient to detect within the micromolar range, and 4) be universally detectable in both HCs and CD individuals, with higher levels in HCs. Based on the first three criteria, we selected four well-characterized meta-organismal metabolites from the Human Metabolome Database (HMDB) [27]: HIPA, phenylacetylglutamine (PAGln), p-cresol sulfate, and phenyl sulfate. Targeted metabolomics of fasting serum from a subset of the treatment-naïve FAH-SYSU IBD cohort (83 HCs and 104 CD) revealed that only HIPA met criterion 4 and was significantly elevated in HCs. By contrast, PAGln was significantly reduced in CD, while phenyl sulfate and p-cresol sulfate showed no significant differences between HCs and CD individuals (Figure S1A).

Targeted measurements of HIPA in the FAH-SYSU cohort (126 HCs and 136 CD) confirmed that fasting serum HIPA levels were significantly higher in HCs than in CD subjects (Figure 1A). Previous studies have identified dietary phenylalanine and polyphenols as sources of HIPA [28–31], however, it remains unclear which is the primary contributor. To identify primary source of HIPA, we used several approaches. First, we correlated serum HIPA levels with consumption of specific food groups in the treatment-naïve FAH-SYSU cohort. Our analysis revealed a positive correlation between HIPA levels and the long-term dietary intake of polyphenol groups in HCs, including chlorogenic acid, one of the most abundant hydrocinnamic acids (Figure 1B). In parallel, we conducted a diet-alternation experiment in mice. Previous studies demonstrate that phenylalanine is metabolised through reductive and decarboxylation pathways to yield 3-phenylpropionic acid and phenylacetic acid [31,32], which subsequently metabolised by the host to form HIPA and PAGln (phenylacetylglycine, PAGly in mice), respectively (Figure 1C). We reasoned that if dietary phenylalanine is the primary source of HIPA, the levels of HIPA and PAGly should exhibit parallel changes in response to dietary alternations. The animals were fed either a grain-based normal chow diet or a high-protein diet in alternating phases, with blood samples collected at designated time points (Figure 1D). Mice maintained on a normal chow diet throughout the experiment served as controls. After four days on the high-protein diet, mice exhibited elevated plasma PAGly levels and reduced HIPA levels. Upon switching back to the normal chow, HIPA levels increased while PAGly levels decreased (Figure 1D). This pattern was consistently observed during the second round of dietary alternation. The results suggested that dietary source of HIPA is primarily from plant component. In addition, we supplement high protein diet with chlorogenic acid, which is one of the most consumed polyphenol in diet, significantly raised plasma HIPA, whereas PAGly remained the same (Figure 1E). Overall, the results suggested that plant polyphenols are the primary contributor to host circulating HIPA.

**Figure 1.**
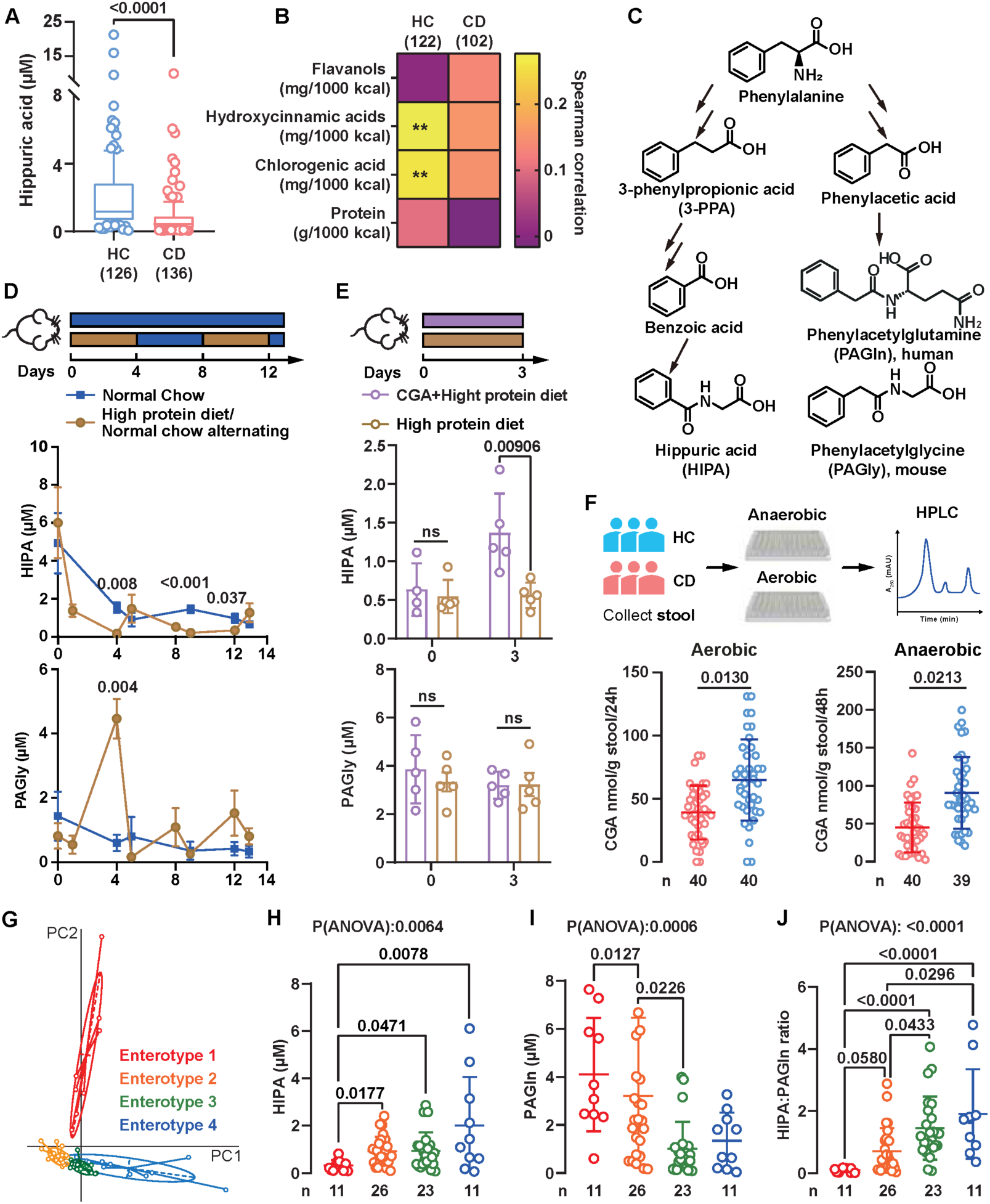
HIPA as a biomarker of dietary polyphenol–gut microbiome interactions. (A) Serum HIPA levels are reduced in CD patients. (B) Correlation of serum HIPA levels with dietary polyphenol intake. (C) Schematic of meta-organismal metabolism showing HIPA and PAGln generation from phenylalanine. (D) Plasma concentrations of HIPA and PAGly in mice following alternation between high-protein and normal chow diets. Significance was determined using multiple t-tests without multiple comparison correction. Mean ± SD; n = 3–5. (E) Plasma HIPA and PAGly levels in mice post chlorogenic acid (CGA) supplementation. Significance was determined using multiple t-tests without multiple comparison correction. Mean ± SD; n = 4-5. (F) CGA degradation potential by faecal microbiome from HCs and CD patients. (G) Gut microbiomes from CD and HCs cluster into four distinct enterotypes. See also Figure S1. (H-J) Serum HIPA, PAGln, and HIPA:PAGln ratio are associated with enterotypes.

### Healthy gut microbiota exhibits greater polyphenol metabolism potential and higher HIPA:PAGln ratio

In addition to dietary polyphenol intake, HIPA concentrations are modulated by gut microbial metabolism. Although the intake of hydrocinnamic acids and chlorogenic acid did not differ significantly between healthy and CD subjects, the positive association between these compounds and circulating HIPA levels observed in HCs was absent in the CD subjects (Figure 1B). We hypothesised that gut microbiota-mediated polyphenol degradation is more efficient in healthy subjects than in individuals with CD. To test the hypothesis, fecal samples from healthy and CD subjects were incubated with chlorogenic acid under both aerobic and anaerobic conditions. Quantification of residual chlorogenic acid revealed significantly lower levels in fecal cultures from healthy subjects relative to CD under both aerobic (24 hours) and anaerobic (48 hours) conditions (Figure 1F), indicating enhanced microbial chlorogenic metabolism in the healthy subjects.

We confirmed that HIPA and PAGln originate primarily from dietary plant and protein components, respectively, and *via* different microbial metabolic pathways. Based on this, we reasoned that fasting serum levels of HIPA and PAGln could serve as functional indicators of gut microbiota structure. Fecal samples from CD patients (n=35) and HCs (n=36) were classified into four enterotypes according to taxonomy composition [33] (Figure 1G). Enterotypes 3 and 4, which were predominantly composed of HCs, exhibited higher levels of HIPA, lower levels of PAGln, and a higher HIPA:PAGln ratio (Figure 1H-J, Figure S1B). In contrast, enterotypes 1 and 2, dominated by CD patients, showed reduced HIPA levels, elevated PAGln levels, and a lower HIPA:PAGln ratio (Figure 1H-J, Figure S1B). Overall, HIPA:PAGln ratio indicates diet-microbiome interaction, with higher HIPA:PAGln ratio indicates microbiome polyphenol degradation efficiency. Interestingly, receiver operating characteristic (ROC) curve analysis revealed that the serum HIPA:PAGln ratio outperformed either metabolite alone in distinguishing patients with CD in the FAH-SYSU cohort, yielding an area under the curve (AUC) of 0.8340, compared to 0.7720 for HIPA and 0.7755 for PAGln (Figure S1C).

### 3-PPA is the predominant physiological precursor of circulating HIPA

The mechanism underlying meta-organismal generation of HIPA remains incompletely understood. HIPA can arise through two pathways, conjugation of benzoic acid with glycine [34], or re-oxidation of 3-PPA mediated by host medium-chain acyl-CoA dehydrogenase (MCAD) [30,35]. To assess the relative contributions of benzoic acid and 3-PPA to circulating HIPA, we analysed plasma samples from mice fed either a custom low-protein diet [36] or standard chow (Figure 2A). We quantified plasma benzoic acid, 3-PPA, and HIPA, along with cinnamoylglycine and phenylpropionylglycine, conjugates of trans-cinnamic acid [30] and 3-PPA [37], respectively. Trans-cinnamic acid, the established precursor of 3-PPA, was also measured [38,39].

**Figure 2.**
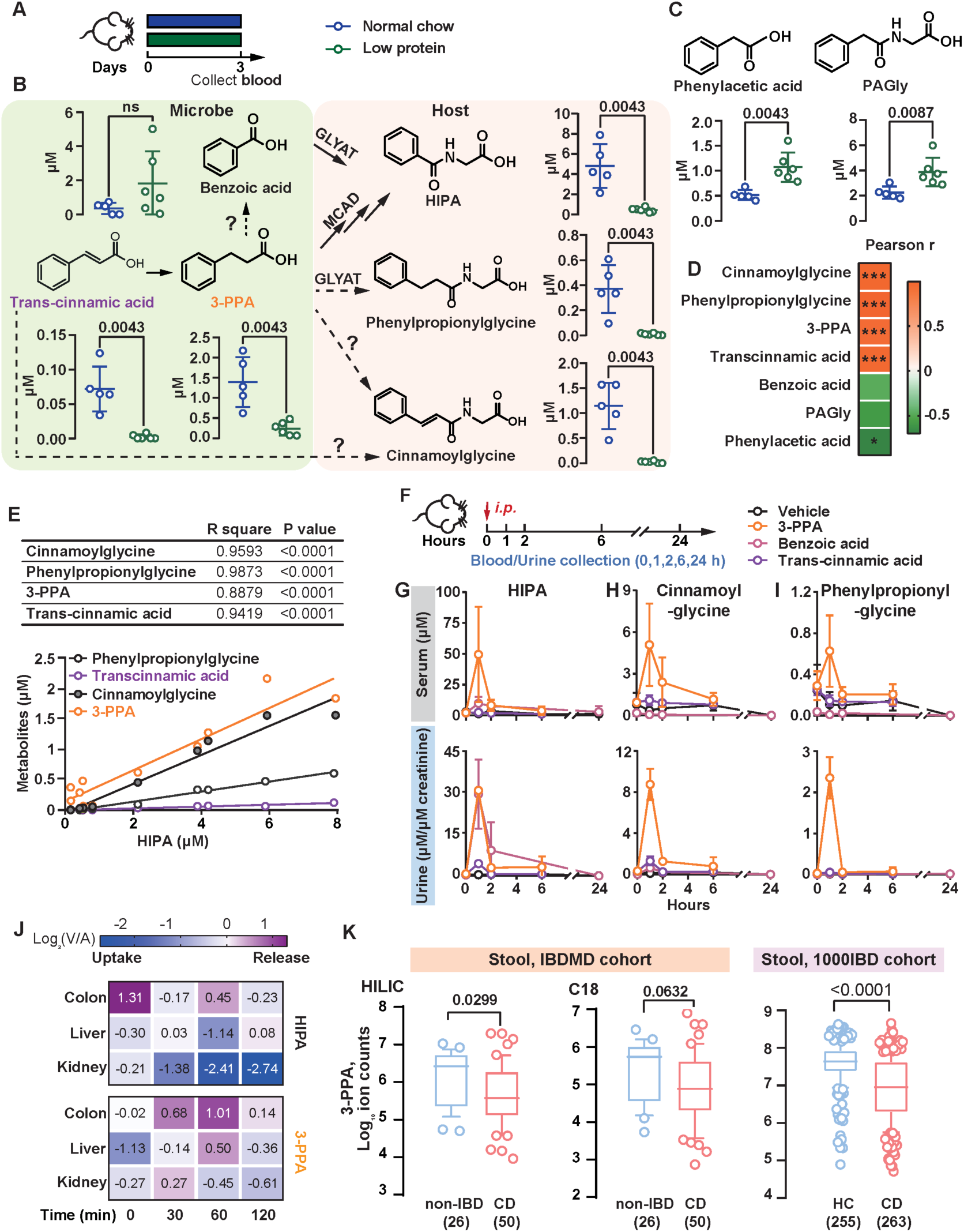
3-PPA is a polyphenol-derived, colon-enriched, microbiome-dependent precursor of HIPA. (A-C) Comparison of plasma HIPA and related metabolites of mice on normal chow and low protein diets. (A) Experimental scheme. (B) HIPA and its related metabolites. (C) Phenylacetic acid and PAGly. MCAD: medium-chain acyl-CoA dehydrogenase. GLYAT: glycine N-acyltransferase. ?: enzyme unidentified. PAGly: phenylacetylglycine. Mean ± SD; n = 5–6. (D, E) The association of HIPA to analysed metabolites in Figure 2B, C. (D) Heat map of HIPA with analysed metabolites. (E) Linear regression of plasma HIPA and positively correlated metabolites. *** P < 0.001, * P < 0.05. (F–I) Systemic levels of HIPA derivatives after *i.p.* injection of 3-PPA, benzoic acid, and trans-cinnamic acid. (F) Study design. Quantification of HIPA (G), cinnamoylglycine (H), and phenylpropionylglycine (I). Mean ± SD; n = 2-10. Only 2 urine samples were collected from trans-cinnamic acid group (0 h) and 3-PPA group (1, 2 h). (J) 3-PPA and HIPA trafficking across organs in LabDiet-fed pigs. Data adapted from Bae et al. (2025) [40]. Log_2_(V/A) represents the log2 ratio of venous (V) to arterial (A) abundances. Data adapted from Bae et al (2025). (K) Stool 3-PPA levels in CD patients compared with non-IBD and healthy controls. Data adapted from Lloyd-Price et al (2019) [4] and Vich Vila et al (2023) [41]. See also Figure S2.

Mice on a chow diet exhibited elevated serum levels of trans-cinnamic acid, 3-PPA, phenylpropionylglycine, and cinnamoylglycine, consistent with elevated HIPA levels, whereas benzoic acid levels did not differ significantly between dietary groups (Figure 2B). We observed that phenylacetic acid and PAGly levels were significantly elevated in mice fed a low-protein diet (Figure 2C). HIPA levels showed positive correlations with cinnamoylglycine, phenylpropionylglycine, 3-PPA, and trans-cinnamic acid, suggesting a shared biosynthetic pathway (Figure 2D, E). No significant association was found between HIPA and benzoic acid levels (Figure 2D).

We next sought to compare the efficiency of host converting benzoic acid and 3-PPA to HIPA. Mice were intraperitoneally injected with 100 mg/kg of either benzoic acid or 3-PPA, and urine and blood samples were collected at specified time points (Figure 2F). 3-PPA and benzoic acid both elevated urinary and plasma HIPA, though 3-PPA drove a stronger plasma-specific response (Figure 2G-I, Figure S2A, B). Unlike benzoic acid, 3-PPA also raised cinnamoylglycine and phenylpropionylglycine levels in plasma and urine, consistent with observations in mice *in vivo* (Figure 2G-I, Figure S2A, B). In humans, 3-PPA is normally metabolised to HIPA *via* MCAD, hence plasma cinnamoylglycine and phenylpropionylglycine levels are low [37]. We also tested trans-cinnamic acid, the precursor of 3-PPA. As expected, trans-cinnamic acid did not elevate HIPA levels in serum or urine (Figure 2G-I, Figure S2A, B). None of the metabolites affected plasma or urine PAGly levels (Figure S2A, B). Overall, the results suggested that 3-PPA is an overlooked direct precursor of HIPA.

### 3-PPA, a diet-modulated and colon-enriched microbiome-derived metabolite, is significantly reduced in CD patients

3-PPA, a diet-dependent microbiome-derived metabolite, reached concentrations about 2 µmol/mg in fresh stool samples from chow-fed mice (Figure S2C-D). Antibiotic treatment (Abx, broad-spectrum cocktail) significantly lowered fecal levels of both trans-cinnamic acid and 3-PPA, along with urinary excretion of HIPA, cinnamoylglycine, and phenylpropionylglycine, confirming their microbial origin (Figure S2C-E). Benzoic acid was not quantified due to fecal matrix effect. Bae et al. (2025) [40] employed a conscious animal model to systematically profile microbe-dependent alterations in xenometabolites and host metabolites. By comparing metabolite abundances in venous (V) and arterial (A) blood, the authors derived organ-specific concentration gradients for each metabolite. Our reanalysis of tissue distribution revealed colon-specific production of 3-PPA, indicated by a positive log₂(V/A) ratio, in contrast to renal clearance of HIPA, indicated by a negative log₂(V/A) ratio (Figure 2J). Both 3-PPA and HIPA were diet-dependent, showing higher levels in control diet–fed pigs compared to those fed a Western diet (Figure S2F, G).

As 3-PPA is a direct precursor of HIPA and produced in the colon, we hypothesised that colonic 3-PPA levels are reduced in CD, consistent with the lower serum HIPA levels observed in CD patients. To test the hypothesis, we analysed publicly available untargeted stool metabolomics data from two independent cohorts, IBDMDB cohort [4] (https://ibdmdb.org/tunnel/public/summary.html) from U.S. and 1000IBD cohort from the University Medical Centre of Groningen, the Netherlands [41]. To minimise longitudinal heterogeneity, only first-visit samples from the IBDMDB cohort were included (n=76). Stool 3-PPA was quantified by HPLC–MS/MS using both HILIC and C18 columns. In this cohort, patients with CD (n=50) exhibited significantly lower 3-PPA levels compared with non-IBD controls (n=26) in HILIC column measurements (P=0.0299), with a consistent but non-significant trend in C18 column analyses (P=0.0632) (Figure 2K). This reduction in 3-PPA was further validated in the 1000IBD cohort, where CD patients (n=263) demonstrated significantly lower stool 3-PPA concentrations compared with HCs (n=255). (Figure 2K).

### 3-PPA and benzoic acid, but not HIPA, ameliorate DSS-induced colitis

Next, we investigated whether 3-PPA supplementation could ameliorate DSS-induced colitis. The downstream metabolite, benzoic acid was included in the investigation, too. SPF mice maintained on a low-protein diet to keep HIPA at low background [36]. Mice received daily *i.p.* injections of either vehicle or test metabolites prior to DSS administration. We monitored body weight and survival over a 9-day experimental period (Figure 3A). DSS treatment induced progressive weight loss, reaching approximately 15% of initial body weight in control mice (Figure 3B). Both 3-PPA and benzoic acid treatment ameliorated colitis compared with vehicle controls, with 3-PPA producing the greatest protection against weight loss (Figure 3B). Intriguingly, while benzoic acid-treated mice showed greater weight loss than those receiving 3-PPA, they demonstrated significantly longer colon lengths (Figure 3C, D), suggesting distinct protective mechanisms. Interestingly, exogenous 3-PPA administration alleviated DSS-induced colitis even in mice on a normal chow diet, despite their significantly higher baseline plasma 3-PPA and HIPA levels compared with low-protein diet mice (Figure S3 A-F). In contrast, HIPA administration (25-100 mg/kg) exacerbated colitis severity, as evidenced by shortened colon lengths and worsened disease activity index (DAI) scores (Figure 3E-G). Pharmacokinetic analysis revealed that plasma HIPA levels peaked rapidly post-injection but returned to baseline within 4 hours due to renal clearance (Figure S3G). This differential response suggests that while HIPA serves as a biomarker of polyphenol-microbiome interactions, 3-PPA and benzoic acids possess direct anti-inflammatory properties.

**Figure 3.**
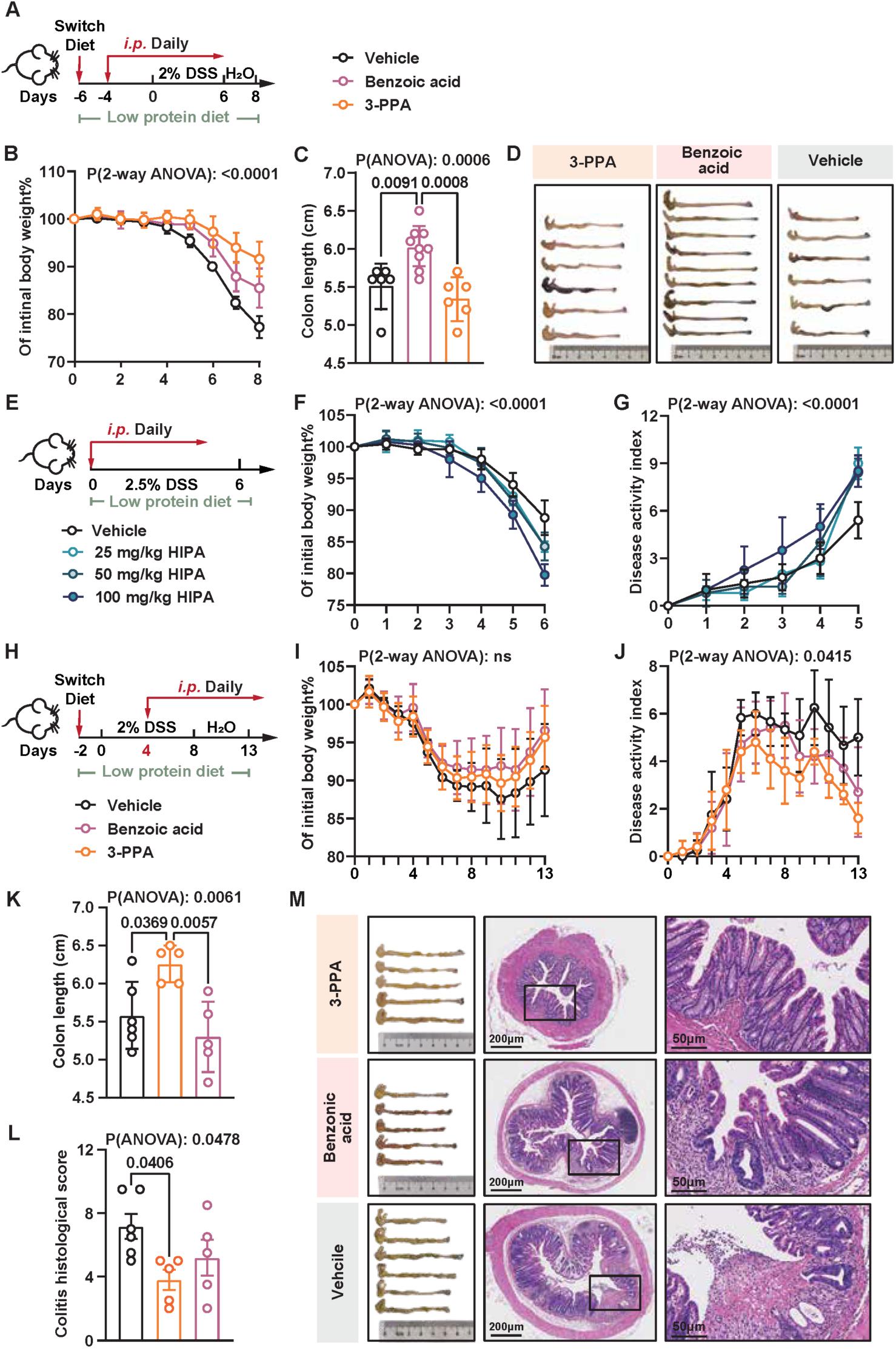
3-PPA and benzoic acid ameliorate DSS-induced colitis. (A–D) Preventive administration of 3-PPA and benzoic acid alleviates DSS-induced colitis, as indicated by improvements in body weight (B) and colon length (C, D). (A) Study design. Mean ± SD; n = 6–9. (E–G) HIPA exacerbates colitis, as shown by greater body weight loss (F) and increased disease activity (G). (E) Study design. Mean ± SD; n = 4–5. (H–M) 3-PPA post-treatment after DSS induction improved colitis more than benzoic acid, with similar body weight recovery (I) but significantly lower disease activity (J), preserved colon length (K), and improved histology (L). Representative colonic morphology and H&E-stained sections at day 13 are shown (M). Mean ± SD; n = 5–6. See also Figure S3.

Beyond the preventive setting, therapeutic intervention initiated 4 days after colitis induction demonstrated that both 3-PPA conferred significant protective effects than benzoic acid (Figure 3H-M). Mice treated with 3-PPA exhibited superior recovery compared with vehicle and benzoic acid groups, including improved DAI scores, preservation of colon length, reduced mucosal erosion, crypt destruction, and inflammatory infiltration, along with a trend toward accelerated body weight regain, although not statistically significant (Figure 3I-M). These findings position 3-PPA as a promising therapeutic metabolite for colitis intervention. We then focused on the anti-inflammatory mechanism of 3-PPA.

### 3-PPA is distinct from its derivatives in acting as a mild PPAR-γ agonist

Phenolic acids act as reactive oxygen species (ROS) scavengers [42]. We first evaluated the effects of various phenolic acids, including 3-PPA, on ROS production of mouse macrophage cell line RAW 264.7 induced by lipopolysaccharide (LPS) stimulation. LPS stimulation led to a significant increase in ROS production, whereas pre-incubation with phenolic acids significantly lowered ROS generation (Figure S4A), however, 3-PPA was equivalent to benzoic acid and caffeic acid.

We reasoned that 3-PPA possesses a unique anti-inflammatory mechanism absent in its derivatives. To investigate this, we compared the colonic tissue transcriptomes of mice treated with 3-PPA versus vehicle (Figure 3A). The analysis revealed enrichment of the peroxisome proliferator-activated receptor (PPAR) signaling pathway (Figure 4A). Specifically, PPAR-γ mRNA expression was significantly downregulated in 3-PPA treated mice, accompanied by significant downregulation of its downstream targets angiopoietin-like protein 4 (*Angptl4)* and perilipin 4 (*Plin4)*, and upregulation of phosphoenolpyruvate carboxykinase 1 (*Pck1)* (Figure S4B).

**Figure 4.**
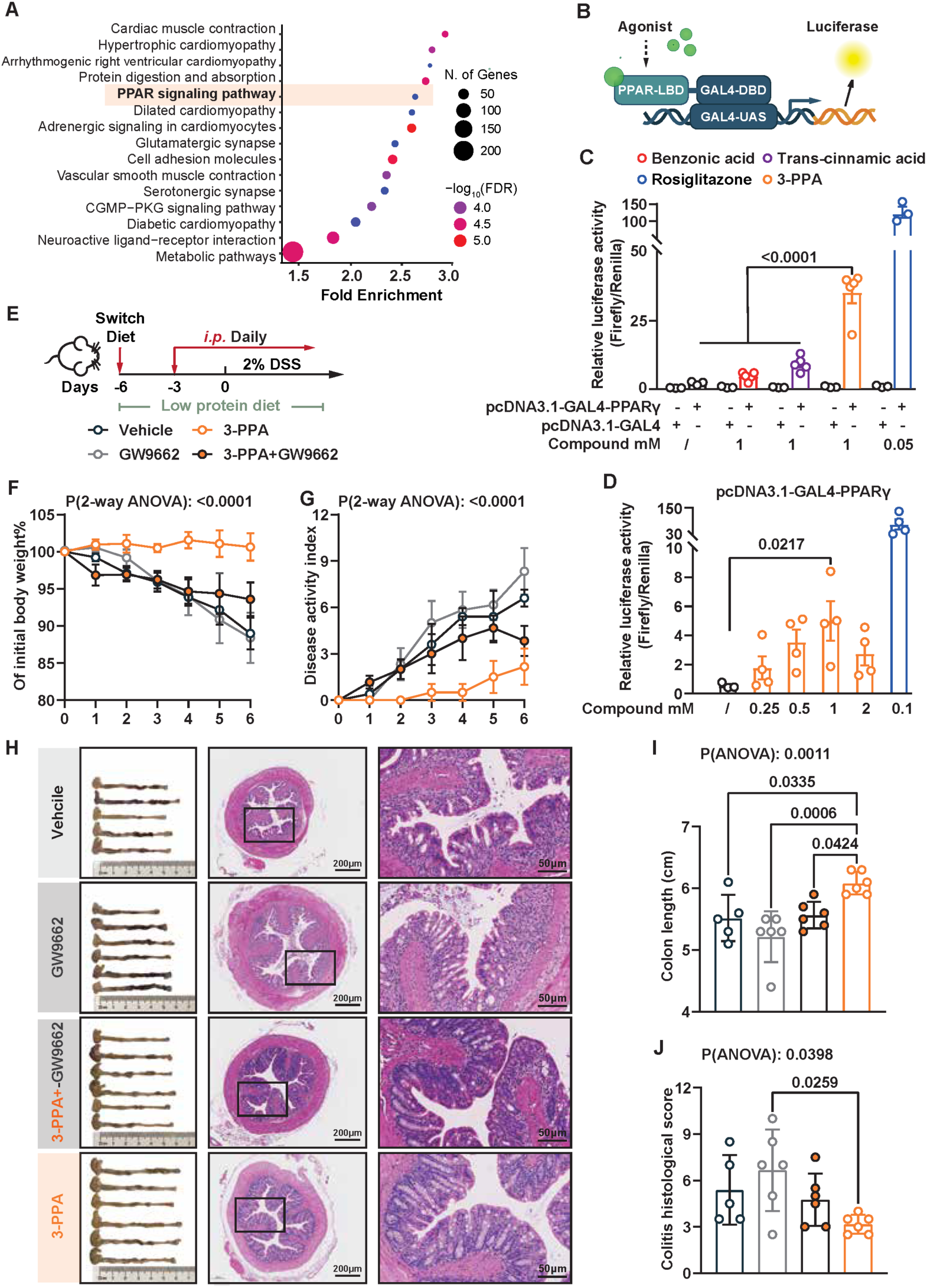
3-PPA ameliorates colitis by activating PPAR-γ. (A) RNA-seq demonstrates PPAR-γ pathway of colonic tissue is enriched by 3-PPA administration from experiment in Figure 3 A. (B) PPAR-γ dual reporter assay scheme for (C, D). (C, D) 3-PPA activates PPAR-γ (C) and exhibits a dose-dependent response (D). One-way ANOVA was performed excluding the rosiglitazone control, with multiple comparisons using Dunnett’s correction. Significant P-values are shown. Mean ± SEM; n = 3-5. (E-J) 3-PPA ameliorates colitis by activating PPAR-γ. (E) Experimental design. Body weight (F) and disease activity (G) were tracked over the duration of the experiment. (H) Colonic morphologies and representative H&E-Stained mouse colon sections at the termination of the experiment on day 6. Colon length (I) and histological assessment of disease severity (J). Mean ± SD; n = 5–6. See also Figure S4.

PPAR-γ has been recognised as a promising therapeutic target for IBD due to its anti-inflammatory and anti-fibrotic properties [14,16]. Prior studies have shown that high concentrations of PPAR-γ agonists, such as rosiglitazone and pioglitazone, can paradoxically suppress PPAR-γ mRNA and protein expression [43,44]. Moreover, 3-PPA derivatives have been reported to act as PPAR-γ agonists [45]. Collectively, these observations led us to hypothesise that 3-PPA itself may function as a PPAR-γ agonist. To test this, we employed a chimeric GAL4–PPAR-γ transactivation assay in HEK293 cells (Figure 4B). In this system, the PPAR-γ ligand-binding domain is fused to the GAL4 DNA-binding domain and activates a GAL4-UAS firefly luciferase reporter upon ligand binding. In the absence of co-transfected GAL4-PPAR-γ, the PPAR-γ agonist rosiglitazone did not increase GAL4-UAS reporter activity, as expected, since reporter activation strictly requires the GAL4-PPAR-γ fusion protein (Figure 4C). In contrast, co-cexpression of GAL4-PPAR-γ conferred robust activation by rosiglitazone (positive control) and revealed that 3-PPA significantly enhanced reporter activity, whereas trans-cinnamic acid showed only minimal activation and benzoic acid had no effect (Figure 4C). 3-PPA reached maximal activation at 1 mM, corresponding to 10–25% of the activity induced by 100 μM and 50 μM rosiglitazone, respectively (Figure 4C, D). Notably, 3-PPA was detected in fecal pellets from SPF mice at ∼2 µmol/µg fresh feces (equivalent to 2 mM; Figure S2B), and in human fecal water at sub-millimolar concentrations [9,10], indicating that the observed bioactivity occurs within a physiologically relevant range.

### 3-PPA ameliorates colitis *via* PPAR-γ *in vivo*

To determine whether 3-PPA mitigates DSS-induced colitis *via* PPAR-γ signaling, we employed the potent, irreversible, PPAR-γ–selective antagonist 2-chloro-5-nitrobenzanilide (GW9662). Mice maintained on a low-protein diet were pretreated with either GW9662 or saline for 3 days prior to DSS administration, followed by daily intraperitoneal injections of 3-PPA (Figure 4E). Administration of 2% DSS in drinking water for 7 days led to significant body weight loss. Compared to vehicle and GW9662-treated controls, 3-PPA significantly reduced weight loss, disease severity, and colon shortening (Figure 4F–J). Co-administration of GW9662 with 3-PPA attenuated these protective effects, resulting in greater weight loss, shorter colon length, and increased disease activity compared to 3-PPA alone (Figure 4F–J), suggesting that PPAR-γ activation contributes to the protective effect of 3-PPA. We also tested rosiglitazone treatment alongside 3-PPA, however, administration of 1 mg/kg rosiglitazone did not ameliorate colitis in our study (Figure S4C–G). Integrating the *in vitro* and *in vivo* findings underscores two key advantages of 3-PPA as a PPAR-γ agonist: 1) it serves as a relatively mild activator; 2) it is a colon-specific agonist, as it is rapidly converted to HIPA systemically and excreted in urine.

### The identified microbial genes responsible for 3-PPA production are enriched in healthy subjects, but low in abundance

We next systematically characterise the gut microbial pathways responsible for 3-PPA biosynthesis. Until now, 4 gene (clusters) have been identified involved in trans-cinnamic acid reduction to 3-PPA. Two genes from *Clostridium sporogenes fldZ* (CLOSPO_02780), which encodes cinnamate reductase [38], and *acdA* (CLOSPO_00312), encoding a 3-(aryl)acrylate reductase [39] (Figure 5A). Another two gene (cluster) are “Flavin” reductase, one is NADH:(hydroxy)cinnamate reductase (*crdAB*) identified in facultative anaerobic marine bacterium *Vibrio ruber* [46], the other one is cinnamate reductase (*cirD*) identified in human associated bacteria *Holdemania filiformis* [47] (Figure 5A).

**Figure 5.**
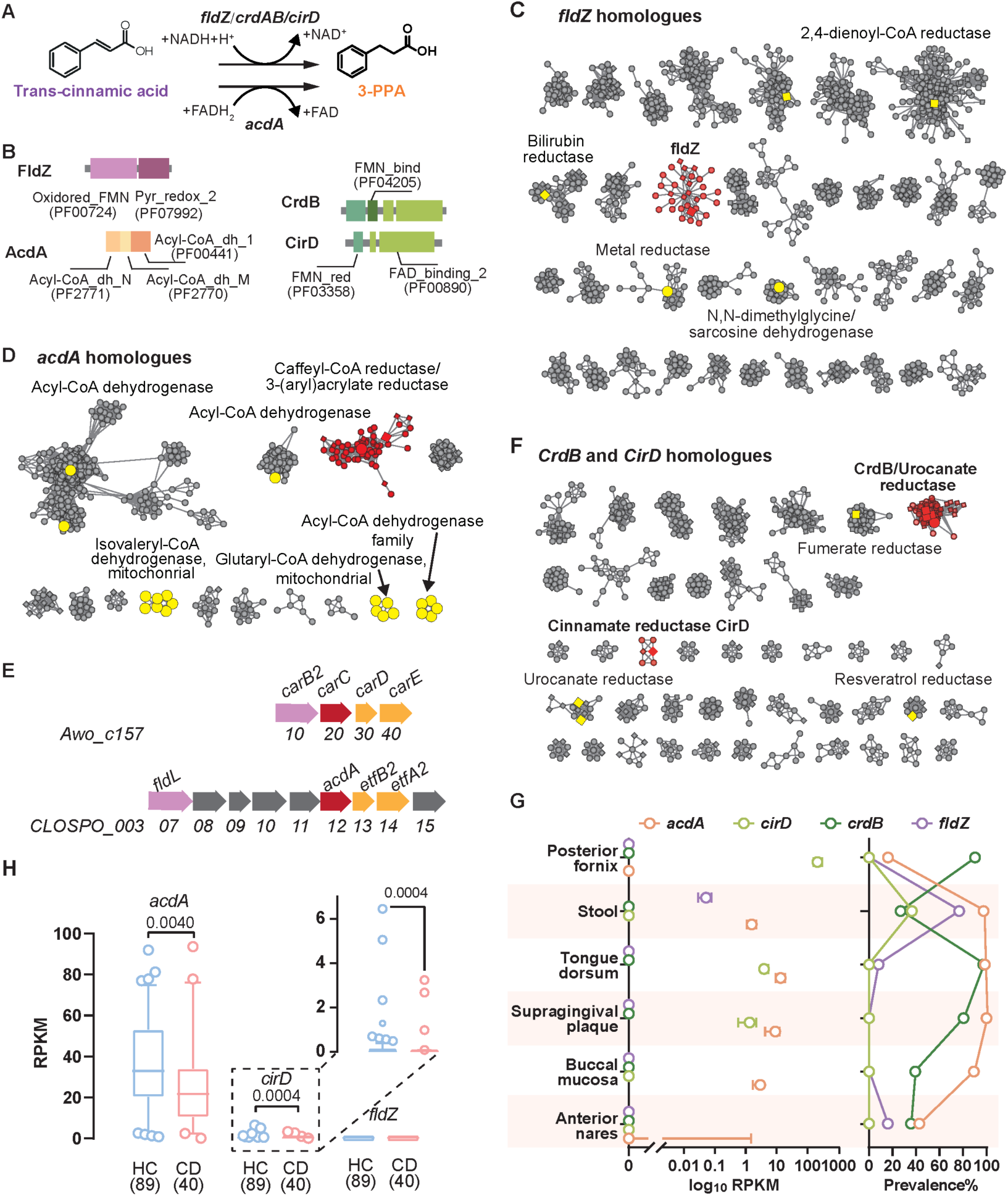
Identified bacterial pathways for 3-PPA production from trans-cinnamic acid are enriched in healthy subjects, with low abundance. (A, B) Identified metabolic pathways and key genes (A) involved in the conversion of trans-cinnamic acid to 3-PPA and their domain structures (B). *fldZ*: cinnamate reductase. *acdA*: 3-(aryl)acrylate reductase. *crdAB*: NADH:(hydroxy)cinnamate reductase subunit A and B. *cirD*: cinnamate reductase. (C, D, F) Sequence similarity networks (SSNs) of *fldZ* (C), *acdA* (D), *cirD* and *crdB* (F). Only partial are shown for clarity. Characterized genes are shown highlighted and enlarged, with those from human-associated microorganisms in diamonds. See also Figures S5–S7 for the complete networks and Supplementary Datasets 1–3 for detailed gene information. (E) Genomic contexts of *acdA* in *C. sporogenes* and *carC* in *A. woodii* ATCC 29683. *carC*, *carD* and *carE* encode Caffeyl-CoA reductase-Etf complex subunits. Light purple: fatty-acyl-CoA synthase. (G) Abundance and distribution of the *fldZ*, *acdA*, *cirD* and *crdB* clusters across 378 HMP metagenomes from six body sites. Median ± SD. See Supplementary Dataset 4 for abundance values. (H) Abundance of *acdA*, *cirD* and *fldZ* in FAH-SYSU metagenomes. Median ± SD. See also Figure S5-8.

To compile a comprehensive list of gut microbial 3-PPA producing gene homologues, we searched the Uniprot protein database using the pfam domains: FldZ (PF00724 and PF07992), AcdA (PF2771, PF2770 and PF00440), and “Flavin” reductase CrdB/CirD (PF00890 and PF03358). These searches yielded 761, 3639 and 1303 unique gene sequences for *fldZ*, *crdB*/*CirD* and *acdA* respectively (Figure 5C-E, Figure S5-7, Supplementary Dataset 1-3). We next classified these homologs using sequence similarity network (SSN) analysis, optimizing the minimum alignment score and amino acid identity thresholds to generate a network in which functionally distinct genes formed separate clusters. We first noticed that the majority of retrieved sequences had unknown functions, and most sequence clusters lacked any characterised members. In the SSN analysis of *fldZ*, *fldZ* formed a distinct cluster separate from 2,4-dienoyl-CoA reductase forms a major cluster, bilirubin reductase, metal reductase, N,N-dimethylglycine/sarcosine dehydrogenase (Figure 5 C, Figure S5, Supplementary Dataset 1). In the SSN analysis of *acdA*, the *acdA* gene in *C. sporogenes* is located adjacent to *carC*, which encodes a caffeyl-CoA reductase in *A. woodii* ATCC 29683 [48] (Figure 5D, Figure S6, Supplementary Dataset 2). *acdA* and *carC* share similar gene cluster structure, each having fatty-acyl-CoA synthase (*fldL* in *C. sporogenes* and *carB2* in *A. woodii*) and electron transfer flavoprotein complex (*etfA2/B2*, *carD*/*E in A. woodii*) in the vicinity (Figure 5E). Indeed, CarC from *A. woodii* has been shown to reduce trans-cinnamic acid to 3-PPA [49]. Beneficial human-associated gut microbes, including *Johnsonella ignava* and *Roseburia inulinivorans* [50], harbor *carC* homologues, suggesting a potential role for *carC* in intestinal 3-PPA production (Figure 5D, Figure S6, Supplementary Dataset 2). Analysis of “Flavin” reductase homologs revealed that CrdB clustered with urocanate reductase, forming a group distinct from CirD, while other characterised reductases, including fumarate reductase, shikimate reductase, and resveratrol reductase, segregated into separate clusters (Figure 5F, Figure S7, supplementary dataset 3).

Using the SSN, we applied ShortBRED to quantify homologues of four 3-PPA–producing genes across 378 high-quality first-visit metagenomes from the HMP, spanning six body sites: stool, buccal mucosa, supragingival plaque, tongue dorsum, anterior nares, and posterior fornix. [51]. *fldZ* and *cirD* clusters prevalence was significantly enriched in stool metagenomics (Figure 5G, Figure S8A, B, Supplementary Dataset 4). Given the phylogenetic and functional similarity between *acdA* and *carC*, the two were combined and collectively referred to as *acdA* in the ShortBRED quantification analysis. *acdA* cluster was detected in stool, with abundance significantly lower than supragingival plaque and tongue dorsum metagenomes (Figure 5G, Figure S8C). The *crdB* cluster is depleted in stool metagenomes, indicating it is not colon-enriched and was excluded from further analysis (Figure 5G).

Next, to assess the DNA levels of for *fldZ*, *acdA* and *cirD* clusters in human gut microbiomes, we used metagenomics data from the FAH-SYSU cohort. We observed that the *fldZ* homologue cluster is dispersed, suggesting it may not represent a functionally uniform group (Figure 5C). Therefore, we compiled 361 sequences from *fldZ* cluster of SSN and constructed a phylogenetic tree (Figure S8D, Supplementary Dataset 5). Phylogenetic analysis revealed that the identified *fldZ* is uniquely characterised by the presence of a DNA-binding MerR family transcriptional regulator adjacent to *fldZ* (Supplementary Dataset 5). In contrast, other 2-enoate reductases within the *fldZ* cluster differ in gene cluster structure, indicating potential functional divergence. Consequently, we identified *fldZ* markers based on 28 *fldZ* genes with adjacent MerR regulators using shortBRED. We found that the abundances of *acdA* and *cirD* clusters were significantly enriched in HCs, however at low levels, with RPKM values ranging from 0.02– 92 for *acdA* and 0–6.452 for *cirD* (Figure 5H). *fldZ* was not detected in the FAH-SYSU cohort metagenomes (Figure 5H).

### Bacterial screening for 3-PPA producers

Both HMP and FAH-SYSU metagenomic analyses revealed low abundances of *acdA*, *cirD* and *fldZ*, despite the high concentrations of stool 3-PPA and urinary/serum HIPA (Figure 5G, H). This suggests the presence of yet unidentified 3-PPA–producing bacteria. Given its health-promoting potential, we aim to screen additional 3-PPA bacteria. 14 dominant commensal bacteria and commercially available probiotic strains were selected and cultured with or without a polyphenol mixture representing commonly consumed polyphenols in the Southern Chinese diet and commercially available, including myricetin, quinic acid, quercetin, rutin, apigenin, genistein, naringenin, trans-cinnamic acid, chlorogenic acid, and caffeic acid [25]. *C. sporogenes* was included in the assay as control. We focused on the metabolism of toxic aglycones, as glycoside hydrolases are widely distributed in human fecal communities, whereas aglycones are variably transformed into non-toxic downstream metabolites [26]. Overall, supplementation with a polyphenol mix significantly inhibited bacterial growth. Notably, *Bifidobacterium breve* produced 3-PPA only in the presence of the polyphenol mix, despite inhibited growth (Figure 6A). In contrast, *C. sporogenes* produced 3-PPA without the polyphenol mix and showed a marked reduction in 3-PPA production, likely due to suppressed growth in the presence of polyphenol mix (Figure 6A).

**Figure 6.**
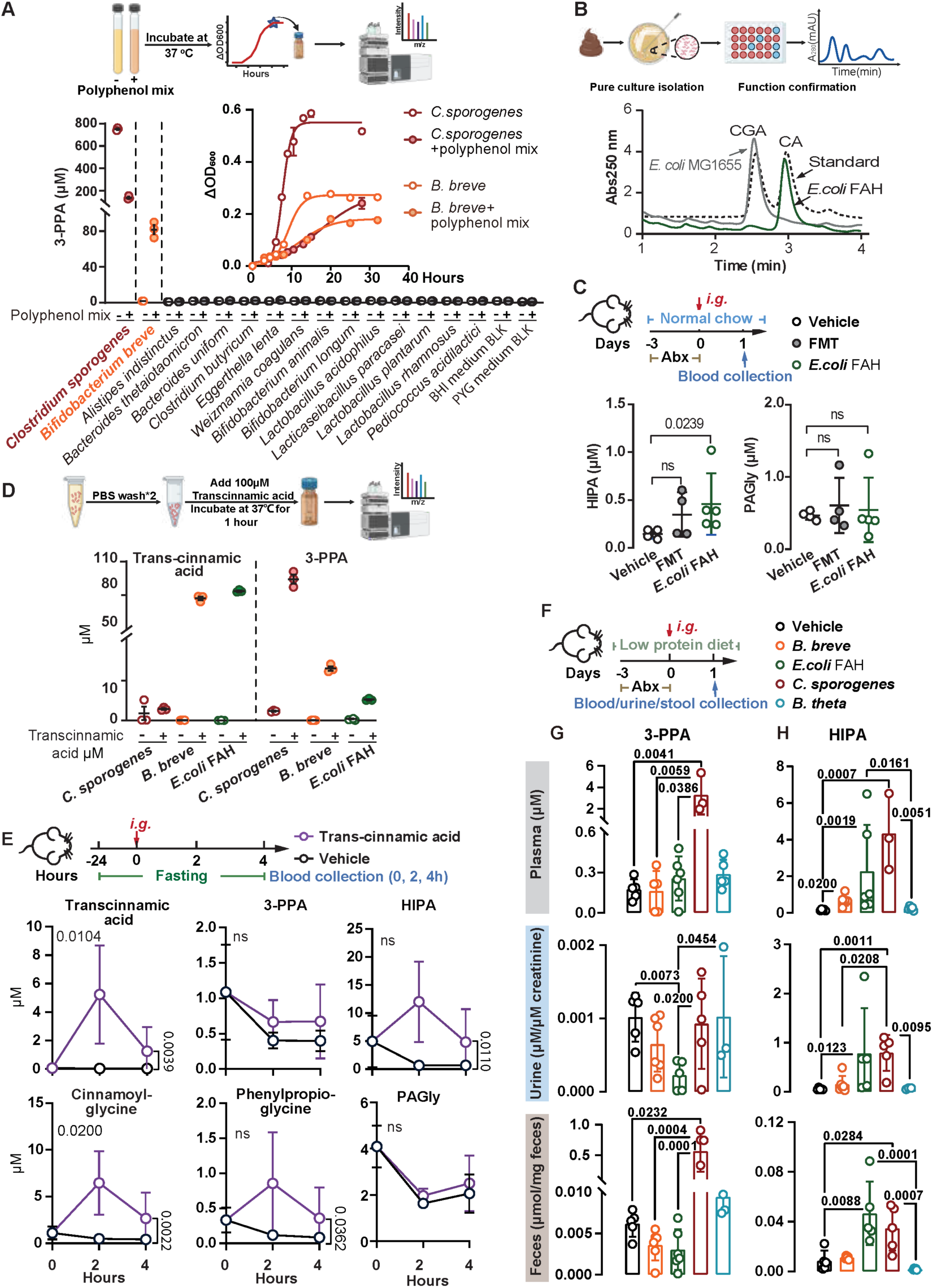
Identification of novel 3-PPA producers. (A) Screening of commensal and probiotic strains for 3-PPA production. Inset: Polyphenol supplementation inhibited bacterial growth. Mean ± SEM. (B) Chromatogram showing healthy subject isolate *E. coli* FAH degraded CGA. (C) Colonisation of *E. coli* FAH raised plasma HIPA levels but not PAGly. FMT: faecal microbiota transplantation using faeces from conventionally raised mice. Mean ± SD; n=4–5. (D) *C. sporogenes*, *B. breve* and *E. coli* FAH converted trans-cinnamic acid to 3-PPA. Mean ± SEM; n=3. (E) Pharmacokinetic profiling of trans-cinnamic acid and related metabolites following oral administration. Mean ± SD; n=3-5. Significance was assessed by two-way ANOVA, with P-values between lines indicating treatment effects. (F-H) Colonization with 3-PPA-producing bacteria increased 3-PPA and HIPA levels *in vivo*. (F) Study design. Plasma, urinary, and faecal concentrations of 3-PPA (G) and HIPA (H) were assessed following bacterial colonization *in vivo*. Significance was assessed using the Kruskal– Wallis test followed by pairwise comparisons without correction for multiple testing. Mean ± SD; n=3–6. See also Figure S9.

A lab isolate, *E. coli* FAH, was obtained from the feces of a healthy donor, whose fecal culture efficiently degraded chlorogenic acid as shown in Figure 1F (Figure 6B). Unlike the reference laboratory strain *E. coli* MG1655, *E. coli* FAH hydrolysed chlorogenic acid to caffeic acid (Figure 6B, Figure S9A, B). Colonization of Abx-treated mice with *E. coli* FAH significantly elevated plasma HIPA, but not PAGly (Figure 6C), suggesting that *E. coli* FAH contributes to 3-PPA production *in vivo*.

We further validated the 3-PPA–producing capacity of these bacteria by incubating cell pellets with trans-cinnamic acid, a direct precursor and a phenolic compound abundant in plants. The results confirmed *C. sporogenes*, *B. breve* and *E. coli* FAH produced 3-PPA in the presence of trans-cinnamic acid (Figure 6D). Neither of *B. breve* nor *E. coli* FAH harbors homologues of *fldZ*, *acdA* or *cirD*, suggesting a different reduction mechanism.

Next, we investigated whether the gut microbiota mediates the *in vivo* conversion of dietary trans-cinnamic acid to HIPA. Oral administration of trans-cinnamic acid significantly elevated plasma HIPA, cinnamoylglycine and phenylpropionylglycine, whereas intraperitoneal administration had no effect (Figure 6E, Figure 2F–I), highlighting the pivotal role of the microbiota in this metabolic pathway. Oral gavage of trans-cinnamic acid resulted in a non-significant trend toward increased plasma 3-PPA, while PAGly levels remained unchanged (Figure 6E).

Given that the gut microbiota mediates trans-cinnamic acid to HIPA metabolism, we reasoned that inoculation with 3-PPA–producing strains would elevate 3-PPA and HIPA levels *in vivo*. We co-administered trans-cinnamic acid with *C. sporogenes*, *B. breve* and *E. coli* FAH. Mice receiving *B. thetaiotaomicron*, a known phenylacetic acid–producing bacterium that generates PAGly *in vivo*, was used as a control (Figure 6F). As expected, colonization with 3-PPA– producing bacteria led to significantly elevated plasma, urinary, and fecal levels of HIPA, along with increased fecal concentrations of 3-PPA, whereas PAGly remained unchanged (Figure 6G–H, Figure S9C, F). Cinnamoylglycine and phenylpropionylglycine exhibited similar upward trends, although some did not reach statistical significance (Figure S9C–E). In contrast, mice inoculated with *B. theta* lead to significantly elevated urine and plasma PAGly levels, while HIPA remained low (Figure 6H, Figure S9F). Examined metabolites remains low in vehicle group without bacteria inoculation, indicating that gut microbiota was not recovered by day 1 following Abx supression (Figure 6F–H, Figure S9C-F). Collectively, these results identify *B. breve* and *E. coli* FAH as novel 3-PPA producers and establish their role in driving the microbiota-dependent conversion of dietary polyphenols into HIPA.

## Discussion

HIPA has been linked to a healthy gut microbiome [52]. In addition to CD, individuals with noncommunicable chronic diseases often exhibit reduced serum levels of HIPA [52,53]. However, the mechanisms governing its production and protective roles remain incompletely understood, limiting its potential as a target for precision nutritional intervention. In this study, we identified the dietary sources, gut microbial metabolic pathways, bioactive intermediates, and physiological benefits associated with HIPA.

First, we confirmed that HIPA serves as a potential functional marker of polyphenol–microbiome interaction. Our findings demonstrate that dietary plant polyphenol intake, in conjunction with microbial production of 3-PPA, synergistically determines circulating HIPA levels. In contrast, protein-derived PAGln reflects microbial metabolism of dietary protein, found elevated in CD, CVD and aging individuals [36,54,55]. Thus, the ratio of HIPA:PAGln ratio represents a metabolic indicator of diet–microbiome interaction. Diet-microbiome-host interactions play crucial roles in regulating human health, yet their direct functional assessment remains challenging. Although shotgun metagenomics provides strain-level resolution and infers functional potential, it is limited in detecting low-abundance species and provides limited insight into active microbial functions [56,57]. What’s more, quantification of dietary intake often relies on subjective questionnaires or food diaries that are prone to recall bias, altered compliance, and insufficient resolution [56]. Thus, metabolomic profiling is advantageous in capturing real-time microbial activity and better reflects diet–microbiome-host interactions. Nevertheless, the potential of HIPA as a functional biomarker of polyphenol–microbiome interactions warrants further validation in real-world cohorts.

Second, we identified 3-PPA as a polyphenol originated, microbiota-derived bioactive metabolite, serves as mild, colon specific PPAR-γ agonist. The intestinal epithelium is in constant contact with a complex microbial environment, and microbial-derived phenolic compounds, such as 4-hydroxyphenylacetic acid [58], urolithin A [59], hydrocaffeic acid [22], have demonstrated a broad spectrum of beneficial properties, including antioxidant, anti-inflammatory activities. Here, we show that 3-PPA is unique as a mild PPAR-γ agonist that ameliorates DSS-induced colitis in mice *via* activation of PPAR-γ signaling. PPAR-γ is known to regulate adipogenesis, inflammation, and apoptosis and has been implicated in the pathophysiology of IBD [11,14,16]. The colon is the primary site of PPAR-γ expression, where epithelial and macrophage PPAR-γ exert protective effects in IBD [14,60]. We therefore used a PPAR-γ antagonist GW9662 to validate its signaling pathway instead of a conditional knockout. Although full PPAR-γ agonists such as rosiglitazone and pioglitazone reduce colitis severity in murine models, their clinical application is limited by adverse effects including weight gain, edema, and heart failure due to systemic full activation [11,19]. We did not reproduce colitis mitigation with 1 mg/kg rosiglitazone reported in other studies, mirroring its relatively narrow therapeutic dosing window.

Consequently, there is growing interest in partial and tissue-selective PPAR-γ agonists as safer alternatives. Our findings suggest that 3-PPA may serve as a naturally occurring, low-affinity, colon specific PPAR-γ modulator, with therapeutic potential. The colon serves as the 3-PPA producing organ, while the kidney functions as the elimination organ for HIPA [40]. Consequently, 3-PPA is efficiently metabolised to HIPA in organs like liver, heart, and kidneys, and is rapidly cleared renally, thus may prevent sustained PPAR-γ activation in extra-colonic tissues. This study highlights the potential of targeting colon-enriched, microbiota-derived polyphenol metabolites as a source of novel bioactive molecules for therapeutic intervention. However, the binding interface of 3-PPA on PPAR-γ and the molecular mechanism underlying its receptor activation warrant further study.

Third, we delineated a diet–microbiome–host pathway for polyphenol metabolism. We observed a strong positive correlation between 3-PPA and circulating HIPA levels, suggesting that systematic HIPA indicate colonic 3-PPA generation. While the *in vivo* pathways mediating 3-PPA formation remain incompletely defined, our study expands the catalogue of microbial enzymes and taxa involved. Beyond previously characterised producers such as *C. sporogenes* and *H. filiformis*, we identified additional contributors, including *carC*, originally implicated in caffeate respiration, which may also catalyse trans-cinnamic acid reduction to 3-PPA. Moreover, we identified *B. breve* and *E. coli* FAH as novel 3-PPA producers, although the underlying molecular mechanisms remain to be elucidated. SSN and gene neighborhood analysis revealed that *bona fide fldZ* is under the control of a MerR-family transcriptional regulator, suggesting that microbial polyphenol metabolism is tightly regulated and responsive to environmental stimulii. Future studies investigating how polyphenols and phenolic acids modulate gene regulation will be critical for elucidating the mechanisms of microbial polyphenol metabolism. Collectively, these results decode functional elements of microbial “dark matter”, uncovering novel biochemical transformations relevant to human physiology. Metagenomic analysis from IBD cohort revealed that HCs show an enrichment of the *acdA*, *cirD*-mediated 3-PPA production pathway. Indeed, human subjects experience significant interindividual variability in their responses to the same food [61]. Furthermore, interindividual variation in polyphenol metabolism suggests that targeted probiotic strategies would optimise the bioavailability of bioactive phenolic acids.

In summary, our work highlights a complex network of interactions among dietary polyphenols, the gut microbiota, and the host immune response and inflammation in CD. Microbiome-derived phenolic acids represent a largely untapped source of colon-specific bioactive molecules with potential health benefits. Our results also suggest that the effects of dietary interventions in CD are not universal. Future precision nutrition strategies should account for individual variations in gut microbiota.

## Limitation of the study

Polyphenols represent a structurally diverse class of compounds, and numerous metabolism pathways and bioactive molecules remain to uncover. The polyphenols tested were selected based on dietary surveys from Southern China. Future studies should consider region-specific differences in polyphenol consumption and metabolism. We examined cross talk between bacteria and host mediated by 3-PPA, however, the inter-bacteria interaction mediated by phenolic acids was not investigated in this study.

## Experimental model and subject details

### Human subjects

All study protocols abided by the Declaration of Helsinki principles and were approved by Ethical Committees of the First Affiliated Hospital of Sun Yat-sen University. Intestinal biopsies and stool specimens were collected as part of the FAH-SYSU cohort study (2016[113]). Subject stool samples were collected at the FAH, SYSU gastroenterology clinic and stored at -80 °C immediately. For culturing assays, fecal samples were collected and diluted to make a 10% (w/v) fecal slurry by resuspension of the feces in 10% (w/v) glycerol solution, and aliquots were stored in cryogenic vials at -80 °C until use. The exclusion criteria applied to all groups were as follows: recent (< 3 months prior) use of any antibiotic therapy, current extreme diet (e.g., parenteral nutrition or macrobiotic diet), known history of malignancy, current consumption of probiotics, any gastrointestinal tract surgery leaving permanent residua (*e.g.*, gastrectomy, bariatric surgery, colectomy), or significant liver, renal, or peptic ulcer disease. Volunteers completed food frequency questionnaires (FFQ) covering 133 food items to assess their usual diet. Dietary polyphenol intake was estimated by a nutritionist using two databases: the China Food Composition Tables, Standard Edition [62], which reports the contents of four polyphenol classes, and the Phenol-Explorer database [63], as described in Xiang et al.[25] [25] Written informed consent was obtained from all participants.

### Mice

Male SPF C57BL/6 mice (6-8 weeks) were maintained on a standard normal rodent diet (Synergy Bio, AIN-93M). All the mice used in this study were bred and raised in the animal facility of the First Affiliated Hospital of Sun Yat-sen University. All animal studies were conducted under protocols approved by the Institutional Animal Care and Use Committee [IACUC] at the First Affiliated Hospital of Sun Yat-sen University (2021 [303], 2021 [035]).

### Cell lines

Human embryonic kidney 293T (HEK293T) and murine macrophage RAW264.7 cells were maintained in DMEM medium supplemented with 10% fetal bovine serum (FBS), 100 U/mL penicillin, and 100 µg/mL streptomycin, at 37 °C in a humidified atmosphere containing 5% CO₂.

### Bacteria

*Bacteroides thetaiotaomicron* ATCC 29148, *Bacteroides uniform* GDMCC 1.898, *Clostridium butyricum* ATCC 19398, *Clostridium sporogenes* GDMCC1.1481, *Alistipes indistinctus* DSM 22530, *Eggerthella lenta* GDMCC 1.990, *Bifidobacterium breve* BBr60, *Weizmannia coagulans* BC99, *Bifidobacterium longum* subsp. *longum* BL21, *Bifidobacterium animalis* subsp. *lactis* BLa80, *Lactobacillus acidophilus* LA85, *Lacticaseibacillus paracasei* LC86, *Lactiplantibacillus plantarum* Lp05, *Lacticaseibacillus rhamnosus* LRa05 and *Pediococcus acidilactici* PA53 strains were routinely cultured in TYG (3% *w/v* tryptone, 2% *w/v* yeast extract, 0.1% *w/v* sodium thioglycolate) or BHI broth at 37 °C in an anaerobic chamber from Coy Laboratories in an atmosphere of 5% H_2_, 10% CO_2_ and 85% N_2_. *Escherichia coli* MG1655 and *Escherichia coli* FAH were routinely cultivated in Luria broth (LB) containing tryptone (10 g·L^−1^), yeast extract (5 g·L^−1^) and NaCl (10 g·L^−1^). Probiotic strains were provided by Wecare Probiotics Co., Ltd. (Suzhou, China).

### Plasmid construction

cDNAs encoding fusion proteins consisting of the N-terminal GAL4 DNA-binding domain (DBD; aa 1-147) and the C-terminal ligand-binding domain (LBD) of human PPAR-γ were synthesised by Dynegene Technologies and subcloned into the multiple cloning site of pcPPAR-γ DNA3.1(+) downstream of the CMV promoter. The resulting constructs were designated pcDNA3.1-GAL4-PPAR-γ. The control plasmid pcDNA3.1-GAL4 encoded only the GAL4 DBD. All plasmids were verified by Sanger sequencing.

## Method details

### Bacteria and cell *in vitro* Experiments

#### Faecal slurry culture

Approximately 20 mg of fecal slurry from HCs and CD patients was inoculated into 150 μL of reinforced medium for clostridia (RMC) supplemented with 150 μM chlorogenic acid in 96-well plates, and incubated anaerobically at 37 °C for 48 h. For aerobic conditions, fecal slurry was cultured in LB broth containing 150 μM chlorogenic acid at 37 °C for 24 h. Following incubation, 20 μL of culture supernatant was mixed with 80 μL of acetonitrile, vortexed for 10 min, and centrifuged (18,000 × g, 20 min, 4 °C). The resulting supernatant was transferred into HPLC vials with inserts and stored at −20 °C until analysis. Chlorogenic acid was quantified using an Agilent 1260 Infinity HPLC system (Agilent Technologies, Santa Clara, CA) equipped with a reverse-phase Kinetex C18 column (50 × 2.1 mm, 2.6 μm, Cat. No. 00B-4462-AN, Phenomenex, Torrance, CA). The mobile phase consisted of solvent A (0.1% acetic acid in water) and solvent B (5 mM ammonium acetate in methanol). Separation was achieved with the following gradient: 0–2 min, 10% B; 2.1–4 min, 10–50% B; 5.1–9 min, 90% B; 9–12 min, 10% B. The flow rate was 0.5 mL/min with an injection volume of 1 μL. Detection was performed at λ = 250 nm using a VWD detector. Each sample was analysed in duplicate, and the mean value was used for plotting.

#### Isolation and whole genome sequencing of *E. coli* FAH

One fecal sample from a healthy subject that exhibited efficient chlorogenic acid degradation was selected for bacterial isolation. The fecal culture was serially diluted and plated on LB and RMC agar plates, followed by aerobic and anaerobic incubation, respectively. Colonies with distinct morphologies were picked and cultured in 96-well plates containing 150 μL RMC or LB medium supplemented with 150 μM chlorogenic acid, and incubated overnight at 37 °C under aerobic and anaerobic conditions. Culture supernatants were processed, and chlorogenic acid levels were quantified as described for **The Fecal Slurry Cultures**. Colonies confirmed to degrade chlorogenic acid were re-streaked 3–5 times to obtain pure cultures, which were then subjected to whole-genome sequencing using Nanopore technology. *E. coli* FAH was deposited at National Center for Biotechnology Information (NCBI: PRJNA923252).

#### Screening for 3-PPA producer

Overnight bacterial cultures were inoculated (1:50, *v/v*) into fresh BHI or PYG medium, with or without a polyphenol mix (100 µM each of myricetin, quinic acid, quercetin, rutin, apigenin, genistein, naringenin, trans-cinnamic acid, chlorogenic acid, and caffeic acid), and incubated at 37 °C. To assess trans-cinnamic acid metabolism, overnight cultures of *C. sporogenes*, *B. breve*, and *E. coli* were subcultured (1:50, *v/v*) into BHI, PYG, and LB media, respectively, and grown to late exponential phase. Cells were harvested by centrifugation at 5,000 × g for 10 min, washed twice with sterile PBS, and resuspended in 1 mL PBS containing either DMSO or trans-cinnamic acid (100 µM) for 3 h. Samples were then collected, centrifuged, and processed for LC-MS/MS analysis.

#### Dual-luciferase reporter assay

To assess PPAR-γ ligand dependent activation, HEK293T cells were seeded in 48-well plates at 5 × 10^4^ cells per well in 200 μL of complete DMEM. After overnight culture, cells were transfected with Renilla luciferase control driven by the SV40 promoter (2 ng/well, internal control) and a firefly luciferase reporter plasmid containing GAL4 response elements (upstream activation sequence, UAS; 180 ng/well), together with either pcDNA3.1-GAL4-PPARγ (30 ng/well) or pcDNA3.1-GAL4 (30 ng/well), using the PEI-based transfection reagent Transporter™ 5 (Polysciences; DNA:PEI ratio 1:3), following the manufacturer’s instructions. The next day, the medium was replaced with fresh complete DMEM (200 μL/well) containing the test compounds: 3-PPA (0.25, 0.5, 1, or 2 mM), benzoic acid (1 mM), trans-cinnamic acid (1 mM), or rosiglitazone (50 or 100 μM, positive control). After overnight incubation, firefly and Renilla luciferase activities was measured using the Dual-Luciferase Reporter Assay Kit, and detected on a multimode microplate reader. Reporter activity was expressed as the ratio of firefly to Renilla luciferase activity. All transfections were performed in triplicate and repeated at least four times.

#### Measurement of Intracellular ROS

Intracellular ROS production was assessed using the oxidant-sensitive probe 2’,7’-dichlorofluorescein diacetate (DCFH-DA). RAW264.7 macrophages (3 × 10^5^ cells/well in 6-well plates) were cultured for 24 h, then serum-starved in DMEM containing 0.4% FBS for 12 h. Cells were treated with 100 μM 3-PPA, benzoic acid, or caffeic acid for 3 h, followed by stimulation with lipopolysaccharide (LPS, 1 μg/mL) for 4 h at 37 °C. After stimulation, cells were incubated with 20 μM DCFH-DA at 37 °C for 40 min, washed three times with PBS, and immediately imaged using an inverted fluorescence microscope. ROS fluorescence intensity was quantified using ImageJ.

### Animal *in vivo* Experiments

#### Dietary intervention

For the diet alteration experiment, mice (n = 4–5) were provided with a custom high-protein diet *ad libitum* for 4 days, switched to normal chow for 4 days, returned to the high-protein diet for 4 days, and finally switched back to normal chow for 2 days. Control mice (n = 4–5) were maintained on normal chow throughout the experiment. Whole-blood samples were collected at designated time points *via* the saphenous vein (survival collection) into heparinised capillary tubes and stored for HPLC-MS/MS analysis.

For the chlorogenic acid supplementation experiment, mice (n = 4–5) were adapted to a custom high-protein diet for 3 days prior to treatment. They then received a daily oral gavage of chlorogenic acid (100 mg/kg in saline) for 3 consecutive days. Control mice (n = 4–5) on the same diet received saline by gavage. Whole-blood samples were collected and stored at day 3 as described above.

For the trans-cinnamic acid supplementation experiment, mice (n = 5–6) were maintained on a normal chow diet and fasted overnight. Mice received a single oral gavage of trans-cinnamic acid (200 mg/kg in 90% saline, 10% DMSO). Blood, urine, and feces were collected at 0, 2, and 4 hours post-gavage. Control mice (n=5-6) received vehicle (90% saline, 10% DMSO).

#### Pharmokinetics

All mice (n = 5–6) received a single intraperitoneal (*i.p.*) injection of 100 mg/kg phenolic acid (benzoic acid, 3-PPA, trans-cinnamic acid) or HIPA in a 200 μL volume. Blood and urine were collected at the indicated time points.

#### Mouse colitis model

For the phenolic acid prevention experiment, mice (n = 5–6) were adapted to a custom low-protein diet for 3 days, then received an intraperitoneal (*i.p.*) injection of 100 mg/kg 3-PPA or benzoic acid (dissolved in 150 μL saline containing 1.84% DMSO, 40% PEG-300, and 5% Tween-80) or vehicle (saline containing 1.84% DMSO, 40% PEG-300, and 5% Tween-80) daily for 4 days prior to DSS (2% w/v) administration and continued throughout the experiment. After 6 days of 2% DSS, the solution was replaced with water for 2 days.

For 3-PPA dose experiment, mice (n=5-6) were maintained on normal chow diet, received the 3-PPA (10, 50, 100 mg/kg prepared as described above) or vehicle 2 days prior to DSS (2% w/v) administration and continued throughout the experiment.

For the phenolic acid treatment experiment, mice (n=5-6) were adapted to a low-protein diet for 2 days before receiving 2% DSS. By day 4, mice exhibited weight loss and soft, bloody stools. Mice were then received an intraperitoneal (*i.p.*) injection of 100 mg/kg 3-PPA, benzoic acid, or vehicle (prepared as above) for the remainder of the experiment. Mice remained on 2% DSS for 4 days before switching to water for recovery.

For the hippuric challenge experiment, mice (n=5-6) were adapted to a low-protein diet for 2 days before receiving 2% DSS, followed by an intraperitoneal (*i.p.*) injection of 25, 50, 100 mg/kg HIPA dissolved in saline or vehicle (saline) for 6 days.

For the PPAR-γ pathway verification experiment, mice were on low protein diet. After 3 days, mice were randomly assigned to four groups (n = 5–6 each): 1) vehicle; 2) GW9662 (5 mg/kg, *i.p.*); 3) 3-PPA (100 mg/kg, *i.p.*); and 4) 3-PPA (100 mg/kg, *i.p.*) + GW9662 (5 mg/kg, *i.p.*). Both compounds were prepared in saline containing 1.84% DMSO, 40% PEG-300, 5% Tween-80 (*v/v*). Mice received GW9662 first, followed 30 min later by 3-PPA. Mice were given 3-PPA and GW9662 3 days before the administration of DSS (2%, *w/v*) and continued throughout the experiment. Another group receiving rosiglitazone (1 mg/kg) dissolved in the same vehicle was also included, following the same procedure described above.

#### Murine antibiotics challenge

Mice were randomly divided into two groups. One group (n = 6) received an antibiotic cocktail in drinking water ad libitum for 3 days, while the control group (n = 6) received plain water. Fecal and urine samples were collected on days 0 and 3 for metabolite analysis as described in HPLC-MS/MS section.

#### Bacteria monocolonisation

Mice were randomly allocated into five groups (n=5-6 per group) and received antibiotics cocktail (Abx) previously shown for 3 days to suppress gut microbiota. Mice were subsequently administered *C. sporogenes*, *B. breve* and *E. coli* FAH *via* oral gavage at a dose of 1.0 × 10^9^ cfu/200 µL each accompanied with 200 mg/kg trans-cinnamic acid. Mice gavaged with saline and *B. thetaiotaomicron* were used as controls. Faecal, plasma and urine samples were collected for metabolite analysis as described in HPLC-MS/MS section.

#### Histopathology

For H&E staining, sections were stained with the Hematoxylin-Eosin (HE) Stain Kit (Solarbio, G1120) and scored using a previously published system [64] as follows: crypt architecture 0 (normal) to 3 (severe crypt distortion with loss of entire crypts); degree of inflammatory cell infiltration 0 (normal) to 3 (intensive inflammatory infiltrate); muscle thickening 0 (base of crypt sits on the muscularis mucosae) to 3 (significant muscle thickening present), goblet cell depletion 0 (absent), 1 (present) and crypt abscess 0 (absent), 1 (present). The colitis histological score was the sum of each mouse.

#### RNA sample preparation, sequencing, and data analysis

Total RNA was extracted from colonic tissue using TRIzol^®^ Reagent (Magen, China) according to the manufacturer’s instructions and quantified with a NanoDrop and an Agilent Bioanalyzer 4150 system. Paired-end libraries were prepared from 1 μg total RNA using the ABclonal mRNA-seq Lib Prep Kit (ABclonal, China), following the manufacturer’s protocol. mRNA was purified using oligo(dT) magnetic beads, fragmented, reverse-transcribed into cDNA, and used for library construction. Sequencing was performed on an Illumina NovaSeq 6000 or MGISEQ-T7 platform at Shanghai Applied Protein Technology. Raw data were filtered and aligned to the Ensembl reference genome GRCm38 using HISAT2 (http://daehwankimlab.github.io/hisat2/). Differential expression and KEGG pathway enrichment between vehicle and 3-PPA groups were analysed with DESeq2 through the iDEP 2.0 platform (https://bioinformatics.sdstate.edu/idep/) [65]. Genes with |log_2_FC| > 0.58 and P_adj_ < 0.1 were considered significantly differentially expressed and included in KEGG enrichment analysis.

#### HPLC–MS/MS

High performance liquid chromatography with in-line tandem mass spectrometry (HPLC-MS/MS) was used for the quantification of multiple metabolites in mouse plasma, urine and stool. Samples (10 μL) were mixed with methanol (40 μL), spiked with internal standards, followed by vortexing and centrifugation to precipitate protein. Urine samples were diluted 1:10 with deionised water before processing. Faeces sample (50 mg) were mixed with methanol with internal standard (200 μL), and vortexed until homogeneous, followed by water bath sonication in ice water for 20 minutes, centrifuge to collect the supernatant. The supernatants were then transferred to glass vials with micro-inserts. The analysis of all calibrators and samples were performed on ultra-performance liquid chromatography coupled to electrospray ionization tandem mass spectrometry platform (SCIEX Triple Quad™ 7500 LC-MS/MS System). Samples were injected onto a reverse-phase Kinetex C18 column (50 × 2.1 mm, 2.6 μm, Cat # 00B-4462-AN, Phenomenex, Torrance, CA). Solvent A (0.1% acetic acid in water) and B (0.1% acetic acid in acetonitrile) were run using the following gradient: 0.0 min (10% B); 0.0–2.0 min (10% B); 2.0–6.0 min (10%B→60%B); 6.0–8.0 min (60%B→90%B); 8.0–10 min (90%B); 10.1 min (90% B→10% B); 10.1–15.0 min (10% B) with a flow rate of 0.3 ml/min and a 1 μL injection volume. The multiple reaction monitoring (MRM) and ion mode used to detect specific compounds of interest are listed. 1) 3-PPA: *m/z* 149.07-105.0441, negative. 2) Trans-cinnamic acid: *m/z* 147.05-103.0191, negative. 3) Benzoic acid: *m/z* 121.04-76.9948, negative. 4) HIPA: *m/z* 179.99-104.9748, negative. 5) Phenylacetylglycine *m/z* 193.8→ 76.1, positive. 6) Cinnamoylglycine: *m/z* 204.1-160.0752, negative polarity. 7) Phenylpropionylglycine: *m/z* 206.1-73.9852, negative. 7) Creatinine: *m/z* 114.03→44.0502, positive. The MRM and ion mode used to detect internal standards are listed as below. 1) D5-phenylacetylglutamine: *m/z* 270.1→130.2, positive. 2) D5-HIPA: *m/z* 183.0985-139.1051, negative. 3) D5-Benzoic acid: *m/z* 126.02-82.051, negative. 4) D9-3-PPA: *m/z* 158.12-114.122, negative. 5) D3-creatinine: *m/z* 117.03→47.0517, positive. The following ion source parameters were applied: nebulizing gas flow, 3 l/min; heating gas flow, 10 l/min; interface temperature, 300°C; desolvation line temperature, 250°C; heat block temperature, 400°C; and drying gas flow, 10 L/min. For data analysis, MultiQuant software (ScieX) was used.

### Bioinformatic analysis

#### SSN analysis of FldZ, AcdA and ‘flavin’ superfamily reductase

Homologues of FldZ carrying PF00724 and PF07992, with characterised 2,4-dienoyl-CoA reductases, yielded 25,107 sequences in UniProtKB. For CirD and CrdB carrying PF00890 and PF04205, 3,648 sequences were retrieved. AcdA (PF00441, PF02770 and PF02771) belongs to the large acyl-CoA dehydrogenase family which yielded >281,549 hits in UniProtKB, exceeding the processing capacity of EFI-EST. Therefore, we performed a BLAST search against UniProtRef90 using AcdA as the query. 94 characterised proteins containing PF00441, PF02770, and PF02771 (including eukaryotic mitochondrial acyl-CoA dehydrogenases) were included as controls to aid in cluster classification, yielding 893 sequences in total. These sequences were submitted to EFI-EST[51,66] (http://efi.igb.illinois.edu/efi-est/) for SSN construction. UniProt IDs, annotations, and PFAM domains are listed in Supplementary Datasets 1–3. The final network was downloaded as 40% representative nodes, with nodes corresponding to characterised enzymes enlarged. Networks were visualised and optimised using Cytoscape v3.10.1.

#### Phylogenetic analysis

361 *fldZ* cluster sequences were aligned and a phylogenetic tree was generated using Clustal Omega multiple sequence alignment web tool [67] (https://www.ebi.ac.uk/jdispatcher/msa/clustalo?stype=protein). The phylogenetic tree is visualised using iTOL 7.2.1 (https://itol.embl.de/) [68]. Gene uniprot ID is available in Supplementary Dataset 4.

#### ShortBRED quantification

SSNs generated with the EFI-EST web tool were subsequently submitted to the EFI-CGFP[51,66] web tool to map metagenomic protein abundances to clusters. The workflow identifies sequence markers unique to families in the input SSN using ShortBRED and clusters them at 85% sequence identity with CD-HIT (CD-HIT 85 clusters). Marker abundances are then quantified across metagenomic datasets and mapped back to SSN clusters. For analysis, a reference library of 380 metagenomes from the Human Microbiome Project (HMP) was used. These metagenomes, derived from healthy adult men and women, represent six body sites: stool, buccal mucosa (inner cheek), supragingival plaque (dental plaque), anterior nares (nasal cavity), tongue dorsum, and posterior fornix (vagina).

To profile the abundance of *acdA*, *cirD*, and *fldZ* in metagenomes from the FAH-SYSU dataset (BioProject: PRJNA793776), markers for *acdA* and *cirD* were obtained from EFI-CGFP (https://efi.igb.illinois.edu/efi-cgfp/) analysis, while a marker for *fldZ* was generated using ShortBRED-Identify from 28 verified and putative sequences against UniRef90 (May 2023) with an 85% identity threshold. These markers were applied in ShortBRED-Quantify to assess gene abundance in paired metagenomes, which had undergone quality control with the KneadData workflow (http://huttenhower.sph.harvard.edu/kneaddata). Gene abundance was reported as reads per kilobase million per million reads (RPKM).

#### Enterotyping

Enterotyping was employed as a methodological approach to categorise the analysed samples into distinct clusters predicated upon the inherent similarities within their microbiome profiles, as outlined in the pertinent literature [33]. The input data for enterotyping were derived from the relative family abundances obtained through metagenomics sequencing. Subsequently, sample clustering based on the relative family abundances was carried out utilizing the Jensen-Shannon Divergence (JSD) distance metric in conjunction with the Partitioning Around Medoids (PAM) clustering algorithm. The selection of the optimal number of clusters was determined by evaluating the Calinski–Harabasz (CH) index, and the statistical significance of this optimal clustering configuration was ascertained by a rigorous comparison utilizing the Silhouette coefficient. Visualization of the enterotyping results was achieved through Principal Coordinates Analysis (PCoA).

#### Quantification and statistical analysis

Statistical analyses were performed with Prism v.8.0 (GraphPad). For comparisons between two groups, statistical significance was assessed using an unpaired t-test for sample sizes ≤6, or the nonparametric Mann-Whitney test for sample sizes >6. Multiple group comparisons were made by Kruskal–Wallis test or ANOVA for most of the studies as indicated. In the box-and-whisker plots, boxes denote the interquartile range (25^th^–75^th^ percentiles), the line within the box indicates the median, and whiskers extend to the 10^th^ and 90^th^ percentiles. Each data point denotes individual human subject, animal, or biological replicate. Statistical details for each experiment are stated in the figure legends with number of samples (or animals) “n” shown within the figures.

## Resource availability

### Lead contact

Requests for further information and resources may be directed to and will be fulfilled by lead contact, Yijun Zhu.

### Materials availability

All resources related to this study, including bacterial strains and plasmids, are available from the lead contact upon request.

### Data and code availability

- Raw metagenomic data of the FAH-SYS cohort were deposited in the NCBI public repository (Bioproject #PRJNA793776).
- HMP IBD metabolomic data can be accessed at https://ibdmdb.org/tunnel/public/summary.html.
- Gene expression profiling data by high-throughput sequencing have been deposited in Gene Expression Omnibus accession no. GSE308767.
- The genomes of the bacterial strains were deposited in public repositories as follows: *E. coli* FAH (NCBI: PRJNA923252), *Lactobacillus acidophilus* LA85 (NCBI: PRJNA1328431)*, Lacticaseibacillus paracasei* LC86 (NCBI: PRJNA1330008)*, Lactiplantibacillus plantarum* Lp05 (NCBI: PRJNA1330416)*, Pediococcus acidilactici* PA53 (NCBI: PRJNA1330483), and *Weizmannia coagulans* BC99 (NCBI: PRJNA809837). *Lacticaseibacillus rhamnosus* LRa05 (CNGB: CNP0008083) and *Bifidobacterium longum subsp. longum* BL21 (CNGB: CNP0008081) were deposited in the China National GeneBank.
- This paper does not report original code.

Any additional information required to reanalyse the data reported in this paper is available upon request.

## Consent for publication

Not applicable.

## Authors’ contributions

Y. Z., R.F., P.B. and M.C. designed research; L.X. and S.Z. performed dietary analysis; L.X., W. Luo., W.Z. and W. Lai. performed animal study; W. Luo. and B.P. performed reporter assay; W.Z. and S.F. performed bacteria experiment; X.W. curated cohort data and conducted bioinformatics analysis; X.L. W.Z. and B.F. performed HPLC-MS/MS analysis; R.F. S.C. and Y.Z. wrote the paper. All authors read and approved the final manuscript.

## Acknowledgments

We thank the First Affiliated Hospital of Sun Yat-sen University Mass Spectrometry Core Laboratory, for their assistance with mass spectrometry analysis. We thank the First Affiliated Hospital of Sun Yat-sen University Research Computing for computational resources, maintenance, and support. We thank Dr. Wei Xie and Dr. Lei Wang for their assistance with the reporter assay. We thank Dr Yin Chen, Dr. Jianping Guo, Dr. Yang Bai and Dr. Xiaoguang Lei for insightful discussions. This work is supported by the National Natural Science Foundation of China (82341217, 82370551 to M.C., 32570124 to Y.Z., 82270579 to R. F, 82222014 to S.C.), National Key Research and Development Program (2023YFC2307004 to Y.Z.), Guangxi Natural Science Foundation (2024GXNSFFA010009 to R. F.).

## References

1. Dolinger, M. et al. (2024) Crohn’s disease. Lancet 403, 1177–1191. 10.1016/S0140-6736(23)02586-2

2. Ananthakrishnan, A.N. et al. (2018) Environmental triggers in IBD: a review of progress and evidence. Nature reviews. Gastroenterology & hepatology 15, 39–49. 10.1038/nrgastro.2017.136

3. Adolph, T.E. and Zhang, J. (2022) Diet fuelling inflammatory bowel diseases: preclinical and clinical concepts. Gut 71, 2574–2586. 10.1136/gutjnl-2021-326575

4. Lloyd-Price, J. et al. (2019) Multi-omics of the gut microbial ecosystem in inflammatory bowel diseases. Nature 569, 655–662. 10.1038/s41586-019-1237-9

5. Lu, Y. et al. (2017) Dietary Polyphenols in the Aetiology of Crohn’s Disease and Ulcerative Colitis-A Multicenter European Prospective Cohort Study (EPIC). Inflamm Bowel Dis 23, 2072–2082. 10.1097/MIB.0000000000001108

6. Kolbel, B. et al. (2024) Low Dietary Flavonoid Consumption Is Associated to Severe Inflammatory Bowel Disease. Gastro Hep Adv 3, 31–37. 10.1016/j.gastha.2023.08.015

7. Koppel, N. et al. (2017) Chemical transformation of xenobiotics by the human gut microbiota. Science 356. 10.1126/science.aag2770

8. Cardona, F. et al. (2013) Benefits of polyphenols on gut microbiota and implications in human health. The Journal of nutritional biochemistry 24, 1415–1422. 10.1016/j.jnutbio.2013.05.001

9. Gill, C.I. et al. (2010) Profiling of phenols in human fecal water after raspberry supplementation. Journal of agricultural and food chemistry 58, 10389–10395. 10.1021/jf1017143

10. Jenner, A.M. et al. (2005) Human fecal water content of phenolics: the extent of colonic exposure to aromatic compounds. Free radical biology & medicine 38, 763–772. 10.1016/j.freeradbiomed.2004.11.020

11. Ahmadian, M. et al. (2013) PPARgamma signaling and metabolism: the good, the bad and the future. Nature medicine 19, 557–566. 10.1038/nm.3159

12. Adachi, M. et al. (2006) Peroxisome proliferator activated receptor gamma in colonic epithelial cells protects against experimental inflammatory bowel disease. Gut 55, 1104–1113. 10.1136/gut.2005.081745

13. Desreumaux, P. et al. (2001) Attenuation of colon inflammation through activators of the retinoid X receptor (RXR)/peroxisome proliferator-activated receptor gamma (PPARgamma) heterodimer. A basis for new therapeutic strategies. The Journal of experimental medicine 193, 827–838. 10.1084/jem.193.7.827

14. Dubuquoy, L. et al. (2006) PPARgamma as a new therapeutic target in inflammatory bowel diseases. Gut 55, 1341–1349. 10.1136/gut.2006.093484

15. Rousseaux, C. et al. (2005) Intestinal antiinflammatory effect of 5-aminosalicylic acid is dependent on peroxisome proliferator-activated receptor-gamma. The Journal of experimental medicine 201, 1205–1215. 10.1084/jem.20041948

16. Su, C.G. et al. (1999) A novel therapy for colitis utilizing PPAR-gamma ligands to inhibit the epithelial inflammatory response. The Journal of clinical investigation 104, 383–389. 10.1172/JCI7145

17. Lewis, J.D. et al. (2008) Rosiglitazone for active ulcerative colitis: a randomized placebo-controlled trial. Gastroenterology 134, 688–695. 10.1053/j.gastro.2007.12.012

18. Schaefer, K.L. et al. (2005) Intestinal antiinflammatory effects of thiazolidenedione peroxisome proliferator-activated receptor-gamma ligands on T helper type 1 chemokine regulation include nontranscriptional control mechanisms. Inflammatory bowel diseases 11, 244–252. 10.1097/01.mib.0000160770.94199.9b

19. Rubenstrunk, A. et al. (2007) Safety issues and prospects for future generations of PPAR modulators. Biochimica et biophysica acta 1771, 1065–1081. 10.1016/j.bbalip.2007.02.003

20. Wright, M.B. et al. (2014) Minireview: Challenges and opportunities in development of PPAR agonists. Molecular endocrinology 28, 1756–1768. 10.1210/me.2013-1427

21. Byndloss, M.X. et al. (2017) Microbiota-activated PPAR-gamma signaling inhibits dysbiotic Enterobacteriaceae expansion. Science 357, 570–575. 10.1126/science.aam9949

22. Bae, M. et al. (2024) Metatranscriptomics-guided discovery and characterization of a polyphenol-metabolizing gut microbial enzyme. Cell host & microbe 32, 1887–1896 e1888. 10.1016/j.chom.2024.10.002

23. Li, Q. and Van de Wiele, T. (2023) Gut microbiota as a driver of the interindividual variability of cardiometabolic effects from tea polyphenols. Critical reviews in food science and nutrition 63, 1500–1526. 10.1080/10408398.2021.1965536

24. Gade, A. and Kumar, M.S. (2023) Gut microbial metabolites of dietary polyphenols and their potential role in human health and diseases. Journal of physiology and biochemistry 79, 695–718. 10.1007/s13105-023-00981-1

25. Xiang, L. et al. (2024) Decoding polyphenol metabolism in patients with Crohn’s disease: Insights from diet, gut microbiota, and metabolites. Food Res Int 192, 114852. 10.1016/j.foodres.2024.114852

26. Culp, E.J. et al. (2024) Microbial transformation of dietary xenobiotics shapes gut microbiome composition. Cell 187, 6327–6345 e6320. 10.1016/j.cell.2024.08.038

27. Wishart, D.S. et al. (2022) HMDB 5.0: the Human Metabolome Database for 2022. Nucleic acids research 50, D622–D631. 10.1093/nar/gkab1062

28. Mostafa, H. et al. (2023) Biomarkers of Berry Intake: Systematic Review Update. Journal of agricultural and food chemistry 71, 11789–11805. 10.1021/acs.jafc.3c01142

29. Tosi, N. et al. (2023) Unravelling phenolic metabotypes in the frame of the COMBAT study, a randomized, controlled trial with cranberry supplementation. Food research international 172, 113187. 10.1016/j.foodres.2023.113187

30. Pruss, K.M. et al. (2023) Host-microbe co-metabolism via MCAD generates circulating metabolites including hippuric acid. Nature communications 14, 512. 10.1038/s41467-023-36138-3

31. Guo, C.J. et al. (2019) Depletion of microbiome-derived molecules in the host using Clostridium genetics. Science 366. 10.1126/science.aav1282

32. Zhu, Y. et al. (2023) Two distinct gut microbial pathways contribute to meta-organismal production of phenylacetylglutamine with links to cardiovascular disease. Cell host & microbe 31, 18–32 e19. 10.1016/j.chom.2022.11.015

33. Arumugam, M. et al. (2011) Enterotypes of the human gut microbiome. Nature 473, 174–180. 10.1038/nature09944

34. van der Sluis, R. et al. (2017) New insights into the catalytic mechanism of human glycine N-acyltransferase. Journal of biochemical and molecular toxicology 31. 10.1002/jbt.21963

35. Derks, T.G. et al. (2008) Neonatal screening for medium-chain acyl-CoA dehydrogenase (MCAD) deficiency in The Netherlands: the importance of enzyme analysis to ascertain true MCAD deficiency. Journal of inherited metabolic disease 31, 88–96. 10.1007/s10545-007-0492-3

36. Feng, R. et al. (2023) Gut Microbiome-Generated Phenylacetylglutamine from Dietary Protein is Associated with Crohn’s Disease and Exacerbates Colitis in Mouse Model Possibly via Platelet Activation. J Crohns Colitis 17, 1833–1846. 10.1093/ecco-jcc/jjad098

37. Rinaldo, P. et al. (1990) The enzymatic basis for the dehydrogenation of 3-phenylpropionic acid: in vitro reaction of 3-phenylpropionyl-CoA with various acyl-CoA dehydrogenases. Pediatric research 27, 501–507. 10.1203/00006450-199005000-00017

38. Dickert, S. et al. (2000) The involvement of coenzyme A esters in the dehydration of (R)-phenyllactate to (E)-cinnamate by Clostridium sporogenes. European journal of biochemistry 267, 3874–3884. 10.1046/j.1432-1327.2000.01427.x

39. Dodd, D. et al. (2017) A gut bacterial pathway metabolizes aromatic amino acids into nine circulating metabolites. Nature 551, 648–652. 10.1038/nature24661

40. Bae, H. et al. (2025) Cross-organ metabolite production and consumption in healthy and atherogenic conditions. Cell 188, 4441–4455 e4416. 10.1016/j.cell.2025.05.001

41. Vich Vila, A. et al. (2023) Faecal metabolome and its determinants in inflammatory bowel disease. Gut 72, 1472–1485. 10.1136/gutjnl-2022-328048

42. Beloborodova, N. et al. (2012) Effect of phenolic acids of microbial origin on production of reactive oxygen species in mitochondria and neutrophils. Journal of biomedical science 19, 89. 10.1186/1423-0127-19-89

43. Hauser, S. et al. (2000) Degradation of the peroxisome proliferator-activated receptor gamma is linked to ligand-dependent activation. The Journal of biological chemistry 275, 18527–18533. 10.1074/jbc.M001297200

44. Haakonsson, A.K. et al. (2013) Acute genome-wide effects of rosiglitazone on PPARgamma transcriptional networks in adipocytes. Molecular endocrinology 27, 1536–1549. 10.1210/me.2013-1080

45. Takada, I. and Makishima, M. (2015) PPARgamma ligands and their therapeutic applications: a patent review (2008 - 2014). Expert opinion on therapeutic patents 25, 175–191. 10.1517/13543776.2014.985206

46. Bertsova, Y.V. et al. (2024) A Redox-Regulated, Heterodimeric NADH:cinnamate Reductase in Vibrio ruber. Biochemistry. Biokhimiia 89, 241–256. 10.1134/S0006297924020056

47. Little, A.S. et al. (2024) Dietary- and host-derived metabolites are used by diverse gut bacteria for anaerobic respiration. Nature microbiology 9, 55–69. 10.1038/s41564-023-01560-2

48. Bertsch, J. et al. (2013) An electron-bifurcating caffeyl-CoA reductase. The Journal of biological chemistry 288, 11304–11311. 10.1074/jbc.M112.444919

49. Hess, V. et al. (2011) A caffeyl-coenzyme A synthetase initiates caffeate activation prior to caffeate reduction in the acetogenic bacterium Acetobacterium woodii. Journal of bacteriology 193, 971–978. 10.1128/JB.01126-10

50. Wang, Z. et al. (2023) Gut microbiota, circulating inflammatory markers and metabolites, and carotid artery atherosclerosis in HIV infection. Microbiome 11, 119. 10.1186/s40168-023-01566-2

51. Oberg, N. et al. (2023) EFI-EST, EFI-GNT, and EFI-CGFP: Enzyme Function Initiative (EFI) Web Resource for Genomic Enzymology Tools. Journal of molecular biology 435, 168018. 10.1016/j.jmb.2023.168018

52. Brial, F. et al. (2021) Human and preclinical studies of the host-gut microbiome co-metabolite hippurate as a marker and mediator of metabolic health. Gut 70, 2105–2114. 10.1136/gutjnl-2020-323314

53. Xu, Y.X. et al. (2024) Alistipes indistinctus-derived hippuric acid promotes intestinal urate excretion to alleviate hyperuricemia. Cell host & microbe 32, 366–381 e369. 10.1016/j.chom.2024.02.001

54. Saeedi Saravi, S.S. et al. (2025) Gut microbiota-dependent increase in phenylacetic acid induces endothelial cell senescence during aging. Nature aging 5, 1025–1045. 10.1038/s43587-025-00864-8

55. Nemet, I. et al. (2020) A Cardiovascular Disease-Linked Gut Microbial Metabolite Acts via Adrenergic Receptors. Cell 180, 862–877 e822. 10.1016/j.cell.2020.02.016

56. Valdes-Mas, R. et al. (2025) Metagenome-informed metaproteomics of the human gut microbiome, host, and dietary exposome uncovers signatures of health and inflammatory bowel disease. Cell 188, 1062–1083 e1036. 10.1016/j.cell.2024.12.016

57. Han, S. et al. (2021) A metabolomics pipeline for the mechanistic interrogation of the gut microbiome. Nature 595, 415–420. 10.1038/s41586-021-03707-9

58. Osborn, L.J. et al. (2022) A gut microbial metabolite of dietary polyphenols reverses obesity-driven hepatic steatosis. Proceedings of the National Academy of Sciences of the United States of America 119, e2202934119. 10.1073/pnas.2202934119

59. Ryu, D. et al. (2016) Urolithin A induces mitophagy and prolongs lifespan in C. elegans and increases muscle function in rodents. Nature medicine 22, 879–888. 10.1038/nm.4132

60. Hontecillas, R. et al. (2011) Immunoregulatory mechanisms of macrophage PPAR-gamma in mice with experimental inflammatory bowel disease. Mucosal immunology 4, 304–313. 10.1038/mi.2010.75

61. Guthrie, L. et al. (2022) Impact of a 7-day homogeneous diet on interpersonal variation in human gut microbiomes and metabolomes. Cell host & microbe 30, 863–874 e864. 10.1016/j.chom.2022.05.003

62. Yang, Y. and Wang, Z. (2018) China food composition tables standard edition Peking University Medical Press

63. Neveu, V. et al. (2010) Phenol-Explorer: an online comprehensive database on polyphenol contents in foods. Database : the journal of biological databases and curation 2010, bap024. 10.1093/database/bap024

64. Wirtz, S. et al. (2017) Chemically induced mouse models of acute and chronic intestinal inflammation. Nature protocols 12, 1295–1309. 10.1038/nprot.2017.044

65. Ge, S.X. et al. (2018) iDEP: an integrated web application for differential expression and pathway analysis of RNA-Seq data. BMC bioinformatics 19, 534. 10.1186/s12859-018-2486-6

66. Zallot, R. et al. (2019) The EFI Web Resource for Genomic Enzymology Tools: Leveraging Protein, Genome, and Metagenome Databases to Discover Novel Enzymes and Metabolic Pathways. Biochemistry 58, 4169–4182. 10.1021/acs.biochem.9b00735

67. Madeira, F. et al. (2024) The EMBL-EBI Job Dispatcher sequence analysis tools framework in 2024. Nucleic acids research 52, W521–W525. 10.1093/nar/gkae241

68. Letunic, I. and Bork, P. (2024) Interactive Tree of Life (iTOL) v6: recent updates to the phylogenetic tree display and annotation tool. Nucleic acids research 52, W78–W82. 10.1093/nar/gkae268

