## Supplementary Figures for "3-Phenylpropionic acid, a microbiota-derived polyphenol metabolite linked to hippuric acid, ameliorates colitis as a colon-enriched PPAR-γ agonist"

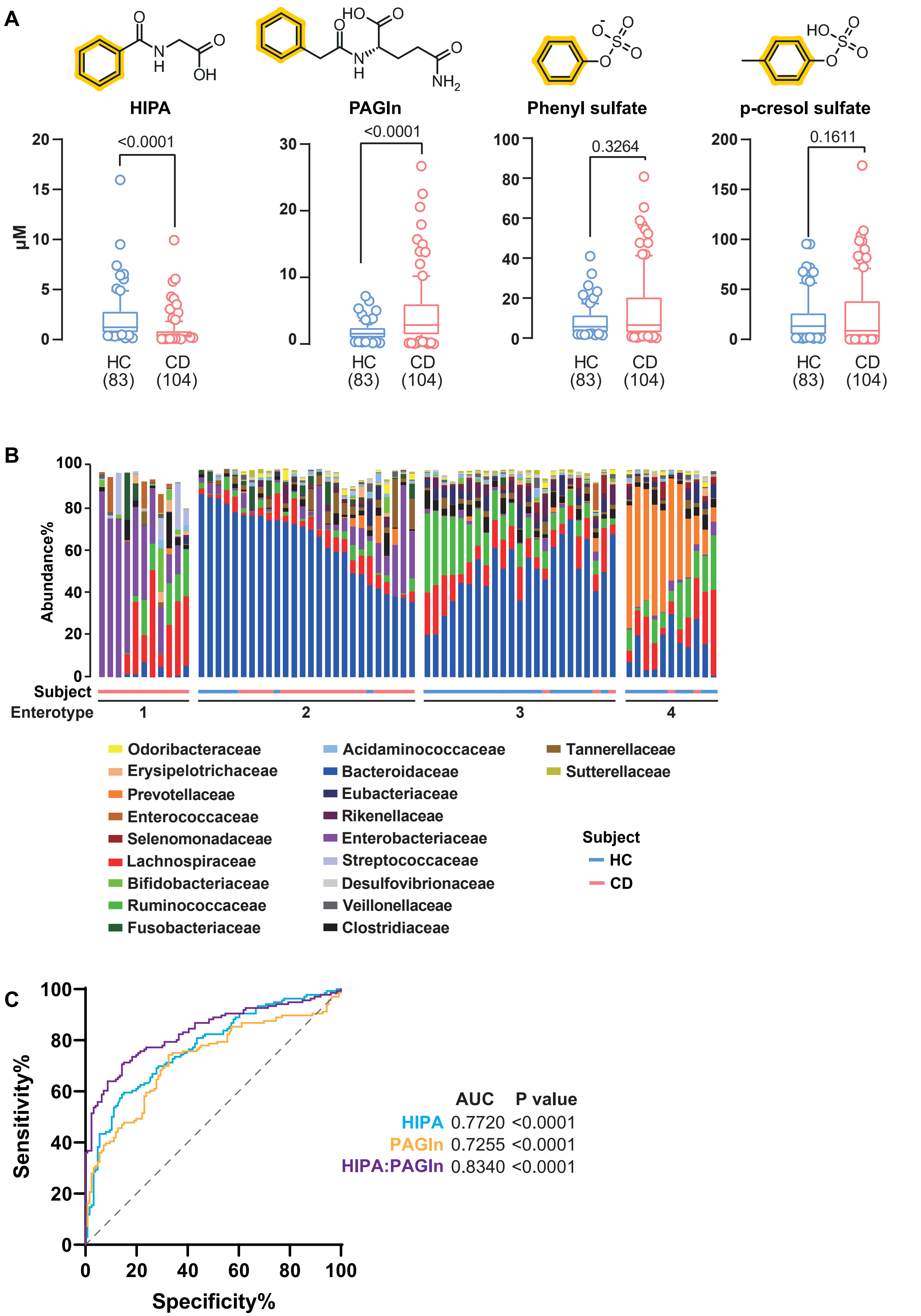
A
HIPA
PAGln
Phenyl sulfate
p-cresol sulfate
μM
<0.0001
<0.0001
0.3264
0.1611
HC
(83)
CD
(104)
B
Abundance%
Subject
Enterotype
1
2
3
4
Odoribacteraceae
Erysipelotrichaceae
Prevotellaceae
Enterococcaceae
Selenomonadaceae
Lachnospiraceae
Bifidobacteriaceae
Ruminococcaceae
Fusobacteriaceae
Acidaminococcaceae
Bacteroidaceae
Eubacteriaceae
Rikenellaceae
Enterobacteriaceae
Streptococcaceae
Desulfovibrionaceae
Veillonellaceae
Clostridiaceae
Tannerellaceae
Sutterellaceae
Subject
HC
CD
C
Sensitivity%
Specificity%
AUC P value
HIPA 0.7720 <0.0001
PAGln 0.7255 <0.0001
HIPA:PAGln 0.8340 <0.0001

**Supplement Fig 2, related to Figure 2**

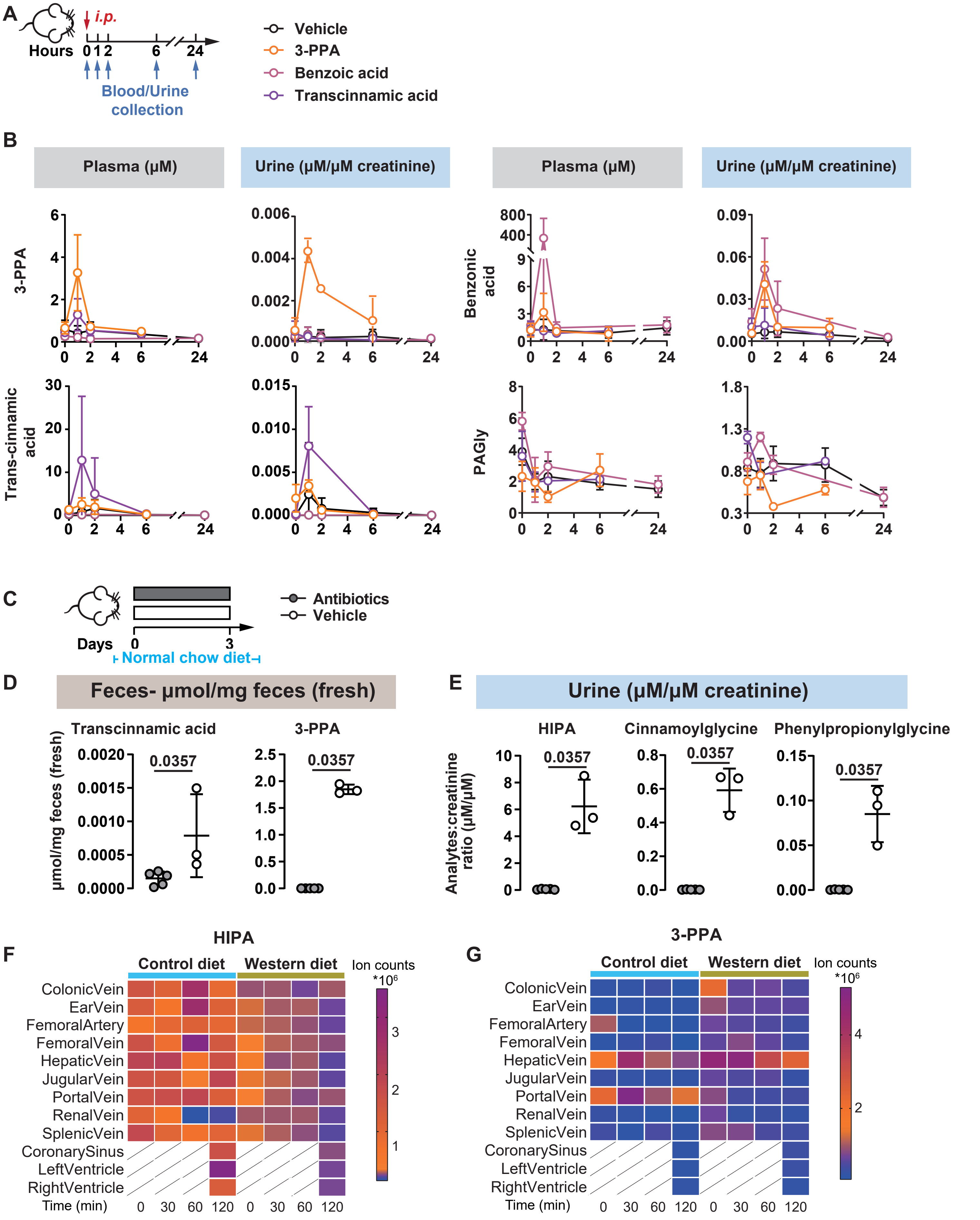

### Supplement Fig 3, related to Figure 3

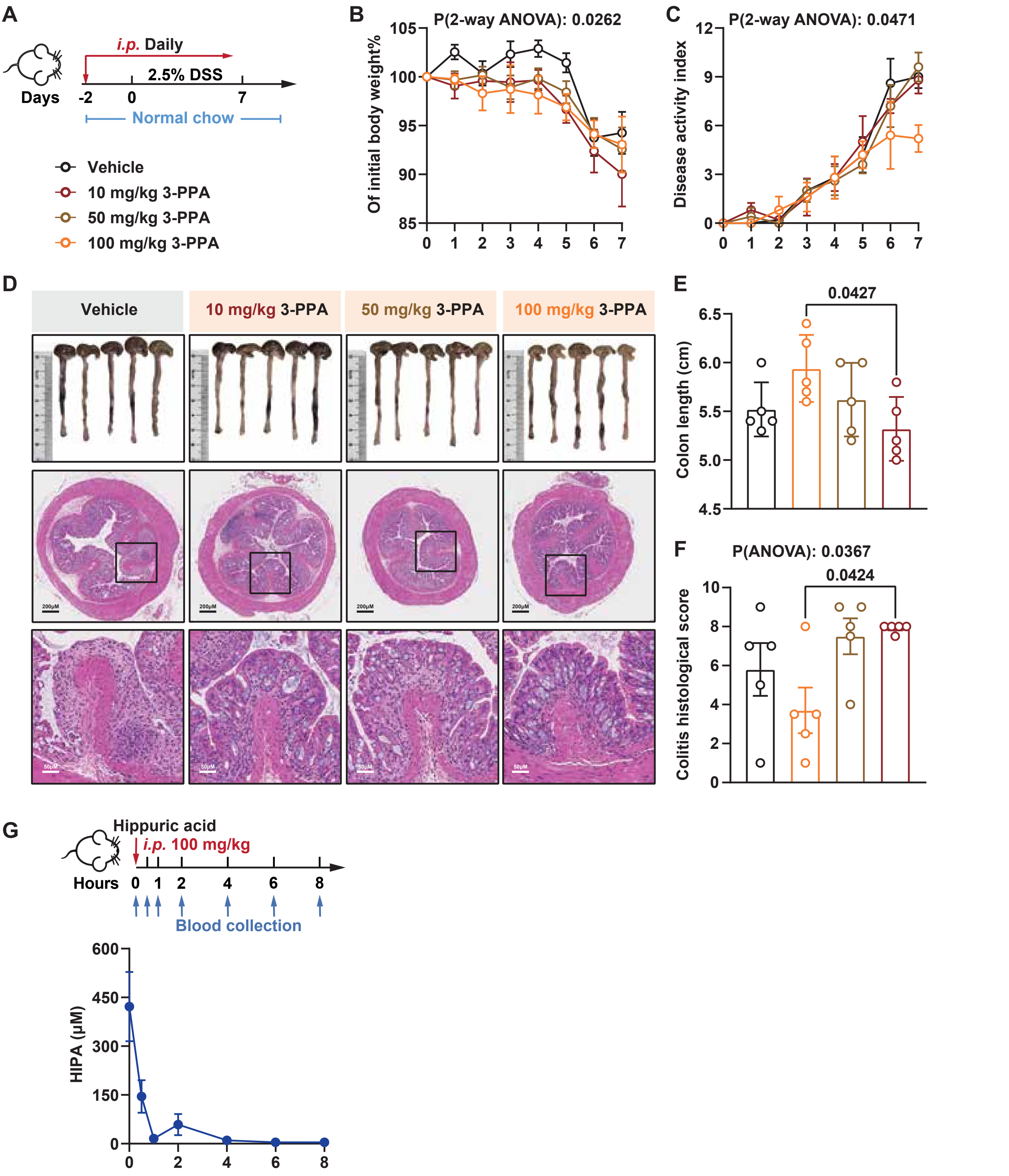

#### Supplement Fig 4, related to Figure 4

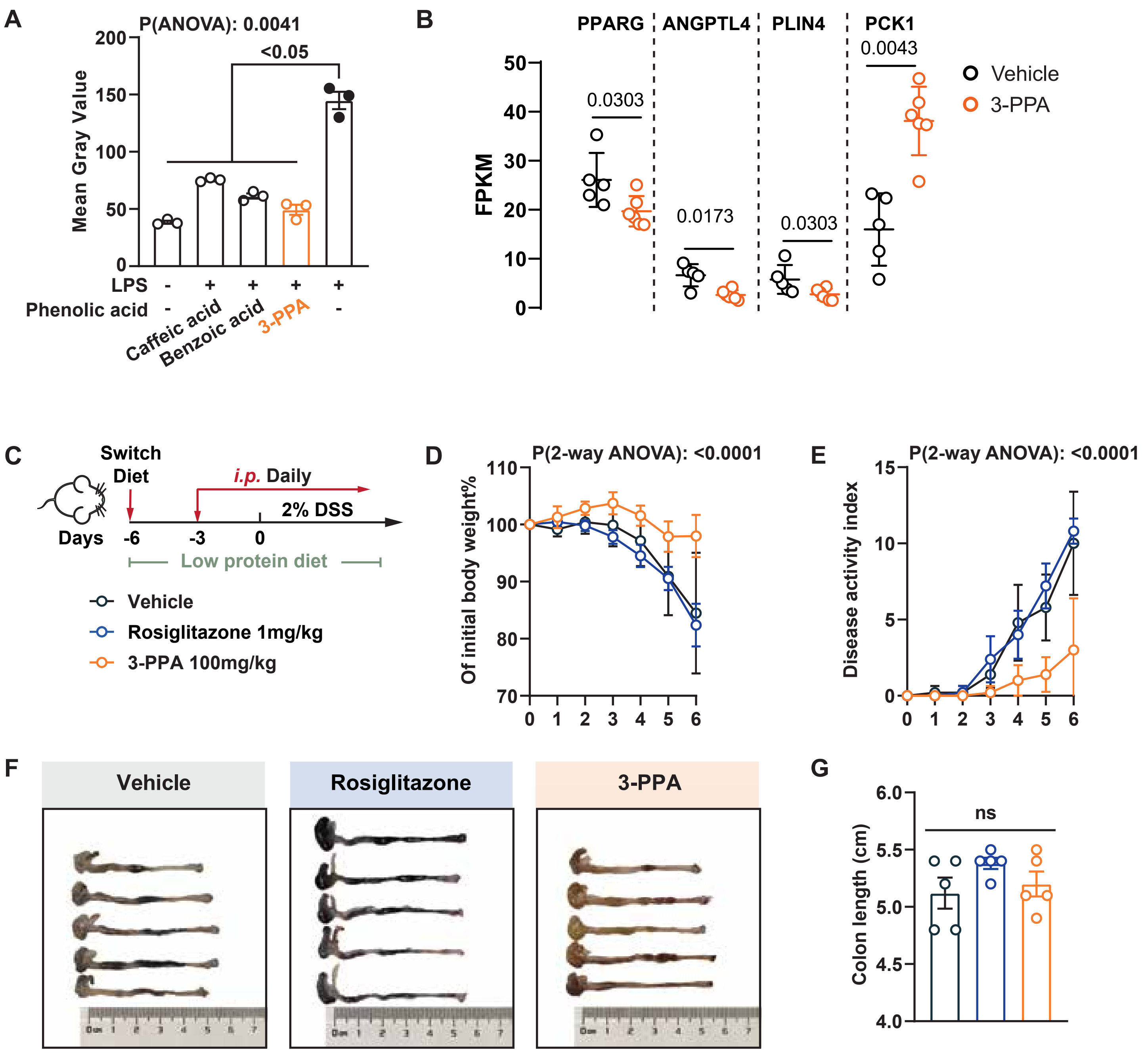

#### Supplement Fig 5, related to Figure 5

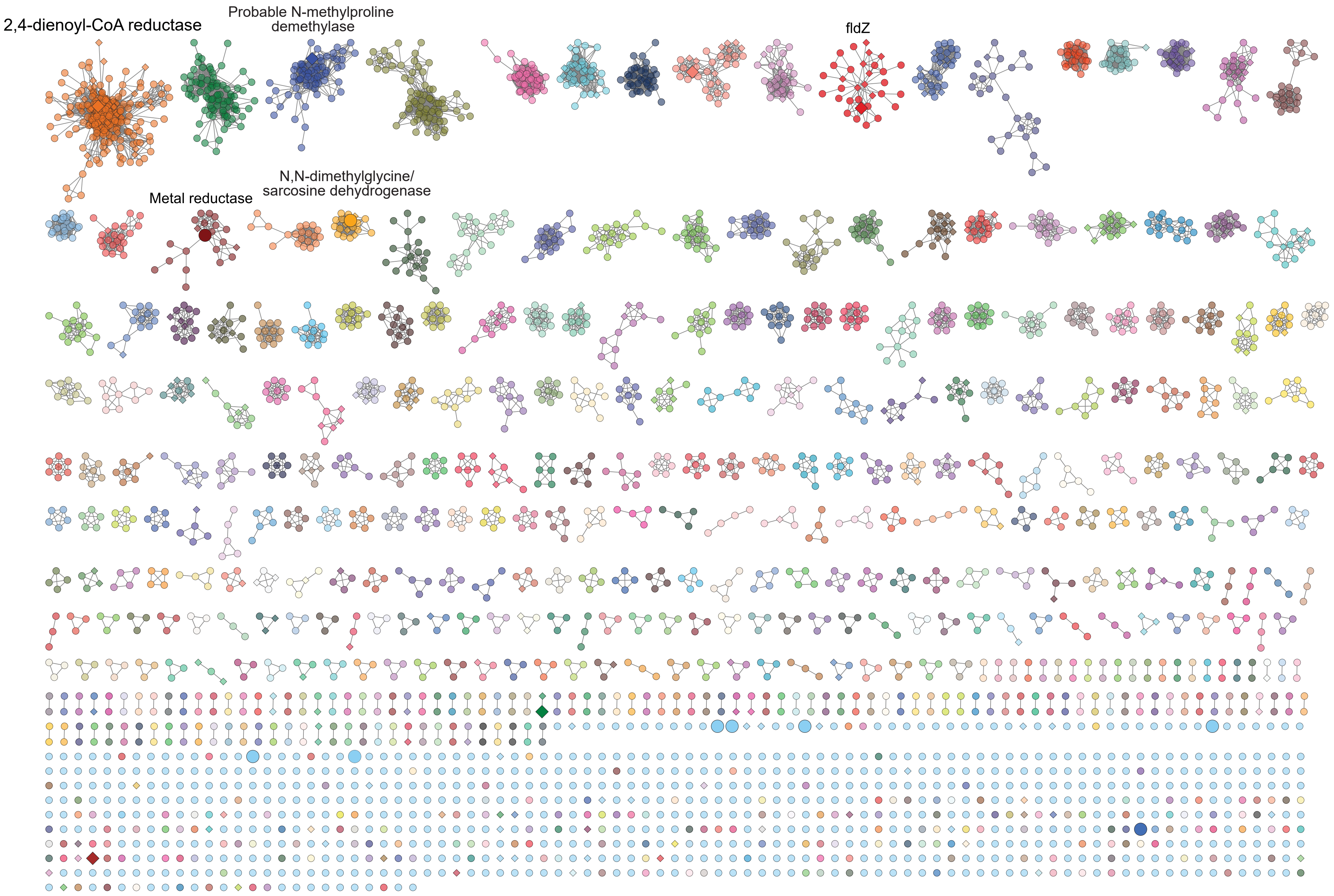

**Supplement Fig 6, related to Figure 5**

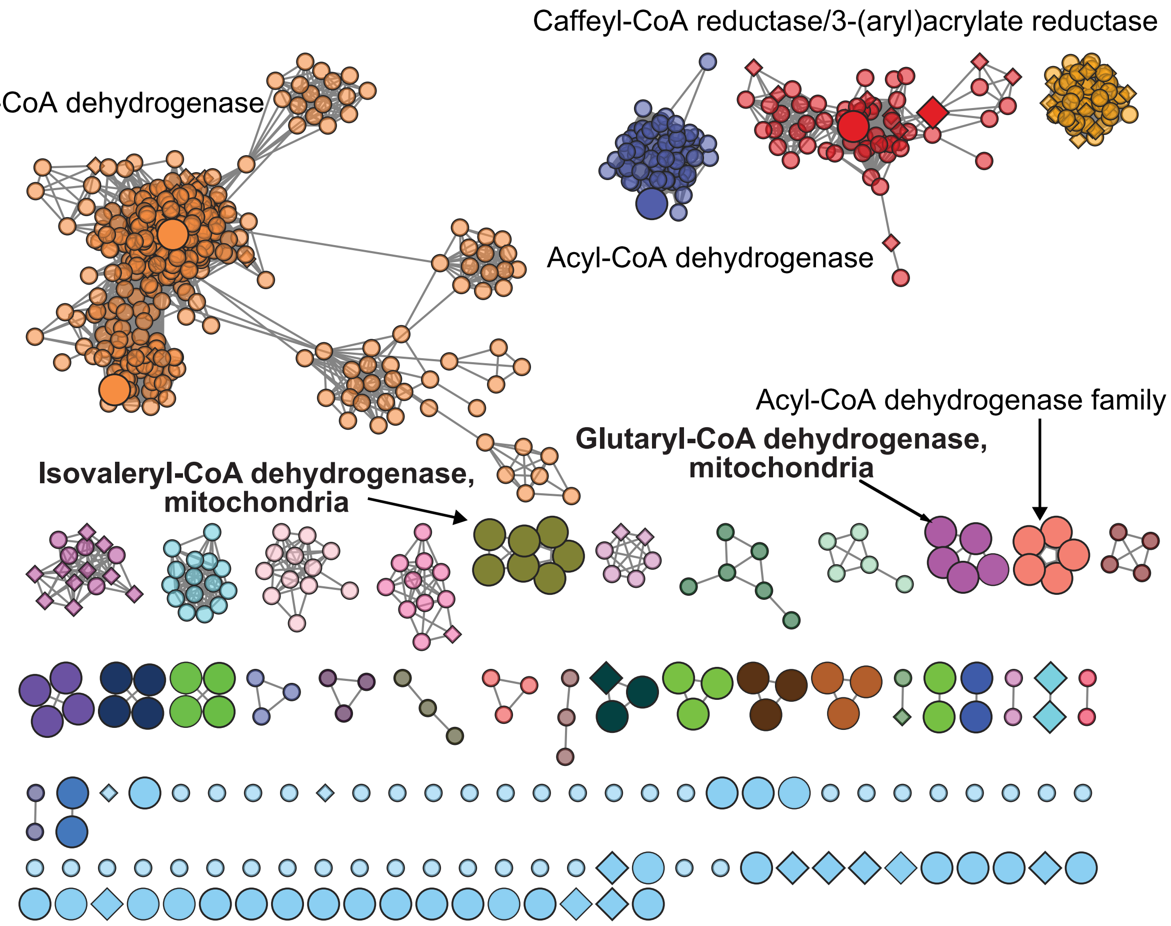

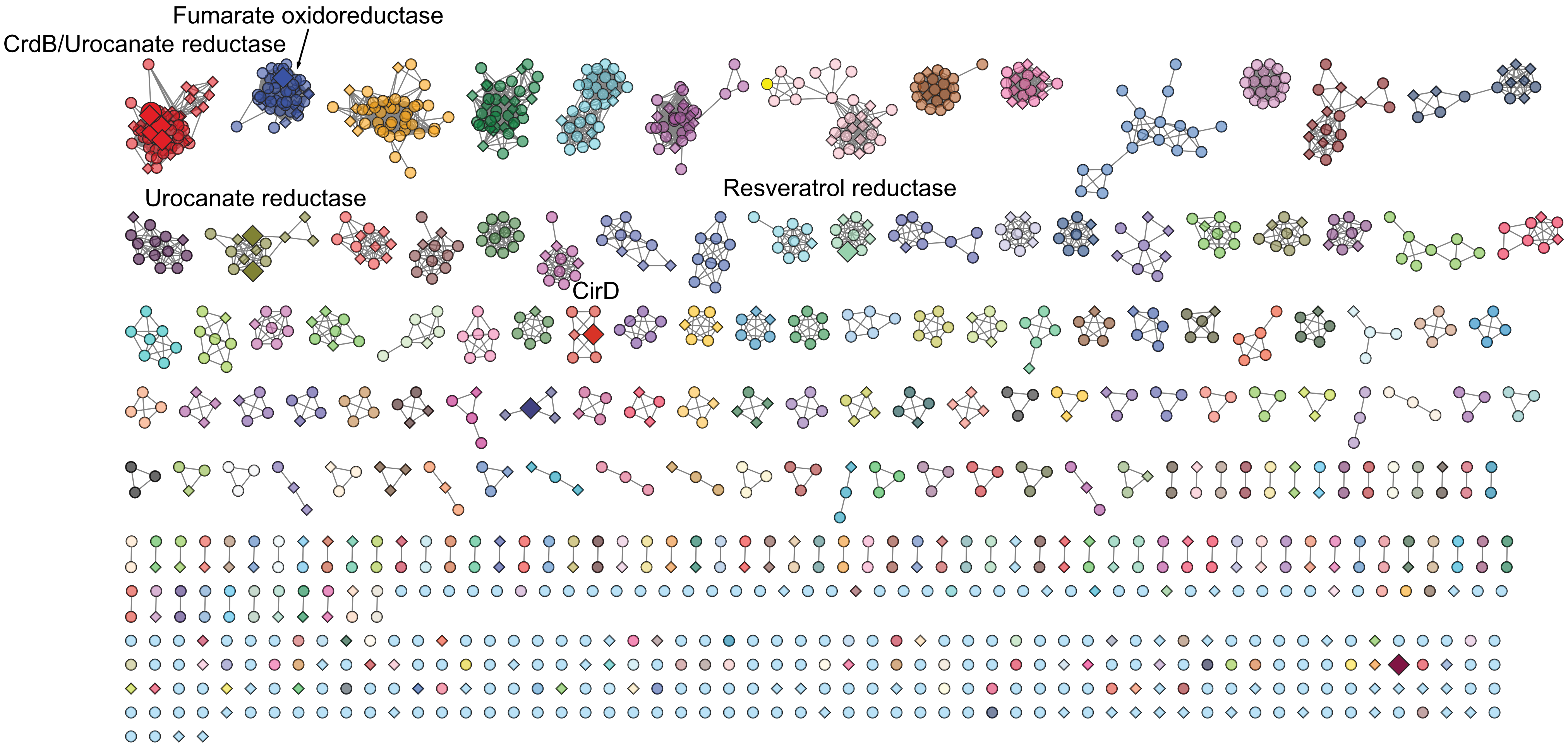
Fumarate oxidoreductase
CrdB/Urocanate reductase
Urocanate reductase
Resveratrol reductase
CirD

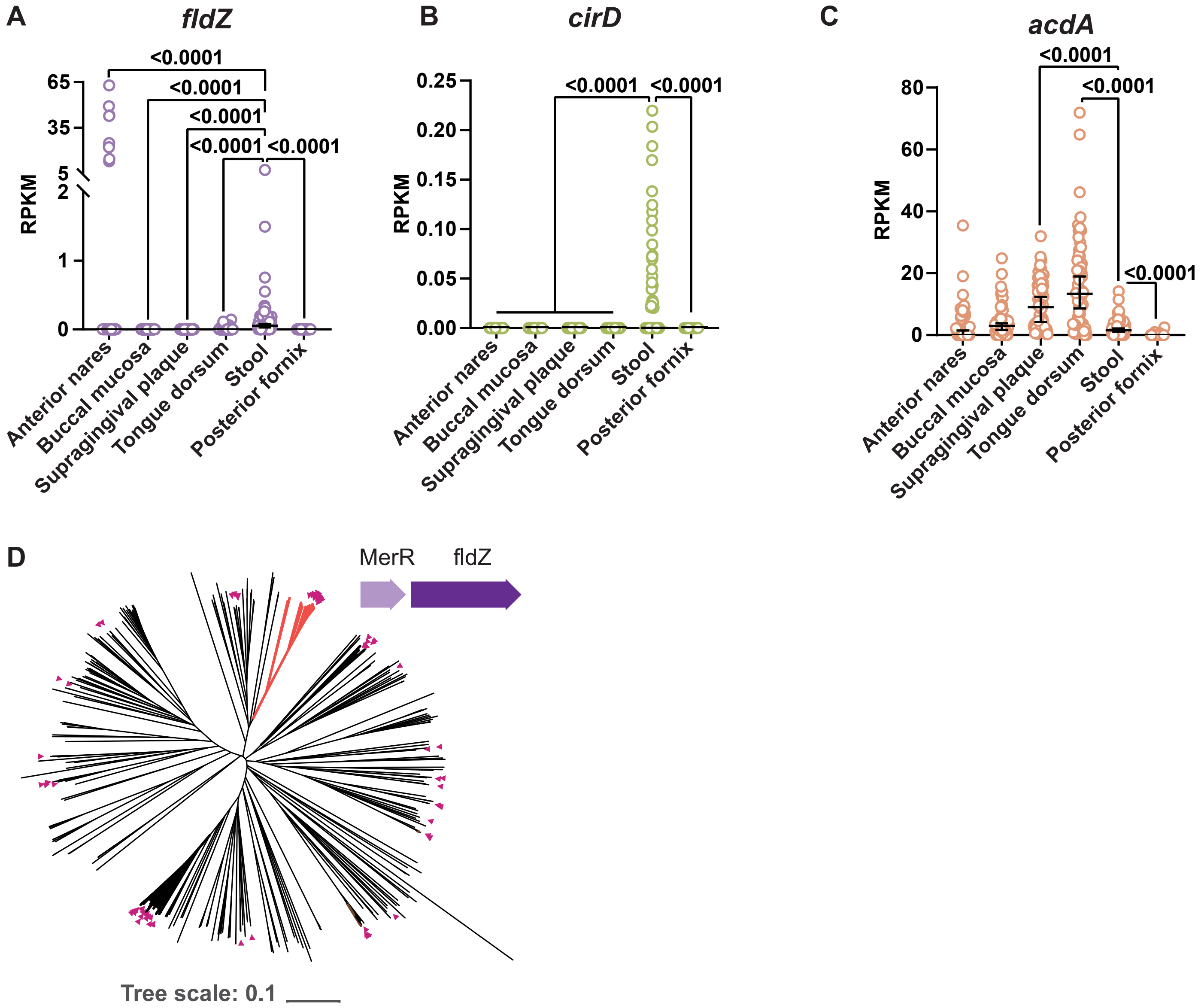
A
fldZ
RPKM
<0.0001
Anterior nares
Buccal mucosa
Supragingival plaque
Tongue dorsum
Stool
Posterior fornix
B
cirD
C
acdA
D
MerR
fldZ
Tree scale: 0.1

**Supplement Fig 9, related to Figure 6**

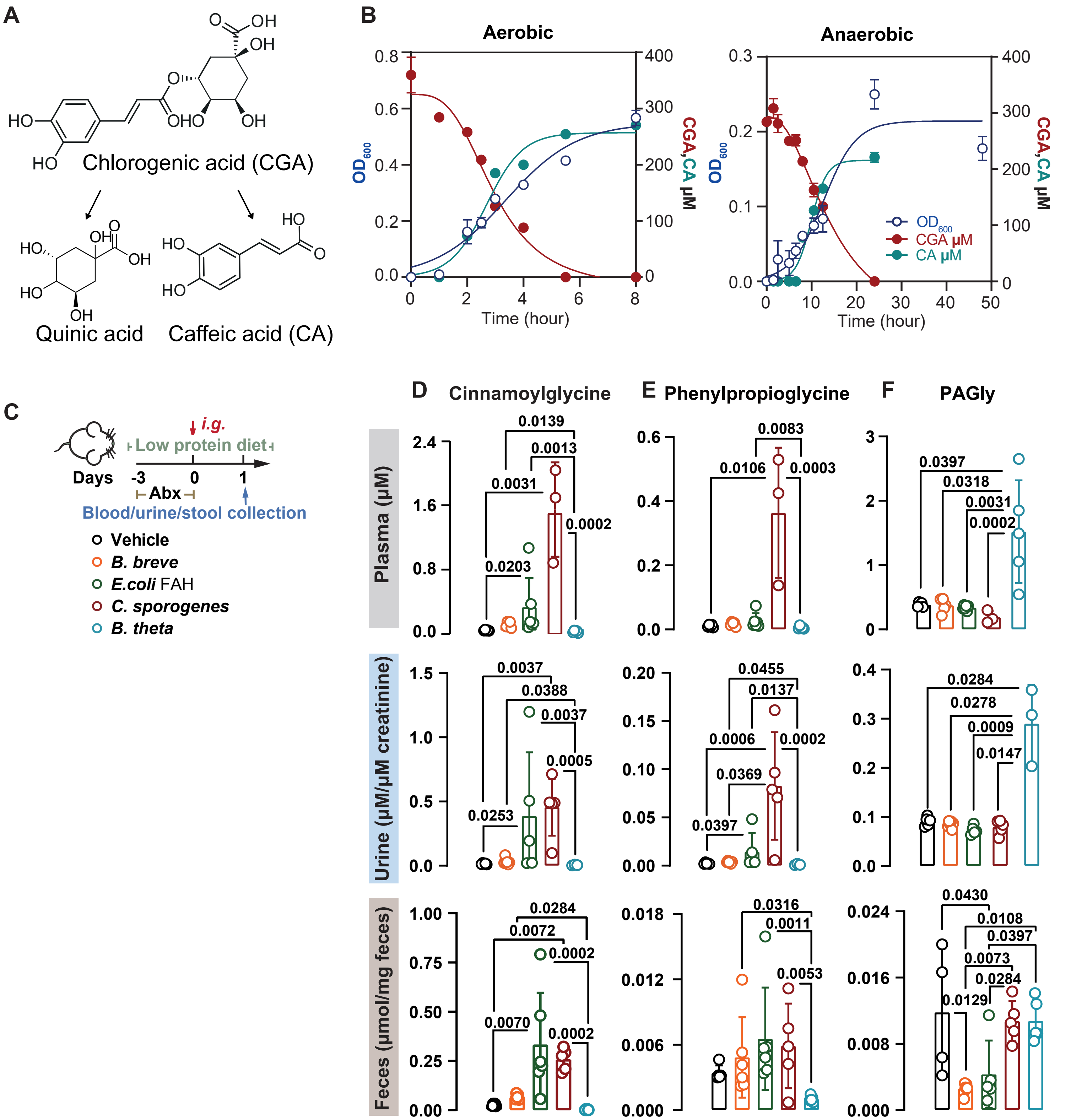
